# Affinity Maturation and Light-Chain-Mediated Paratope Diversification Anticipate Viral Evolution

**DOI:** 10.1101/2025.08.27.672735

**Authors:** John Dingus, Duck-Kyun Yoo, Sachin Kumar, Yajuan Wang, Md Golam Kibria, Shahab Saghei, Zahra Allahyari, Jessica W. Chen, Natalie M. Caputo, Jason Hwang, Bing Chen, Duane R. Wesemann

## Abstract

A key goal of vaccinology is to train the immune system to combat current pathogens while simultaneously preparing it for future evolved variants. Understanding factors contributing to anticipatory breadth, wherein affinity maturation against an ancestral strain yields neutralization capacity against evolved variants, is therefore of great importance. Here, we investigated the mechanism of anticipatory breadth development in a public antibody family targeting the functionally restricted ACE2 binding site on SARS-CoV-2. IGHV3-53/66 antibodies isolated from memory B cells of infection-naïve individuals vaccinated with the ancestral Wuhan-strain mRNA vaccine frequently neutralized evolved Omicron variants and contained several hallmark mutations previously shown to enhance neutralization breadth. Comparative analyses with antibodies from Omicron breakthrough infections revealed that breadth-associated patterns of somatic hypermutation emerged independently of variant exposure. However, Omicron infection had a marked impact on light chain pairing frequencies, suggestive of variant-imposed selection of favorable light chains. Analysis of available IGHV3-53/66 antibody structures complexed with SARS-CoV-2 receptor binding domain (RBD) clarified these findings; convergent somatic mutations on the heavy chain largely refined contacts with invariant RBD residues, while light chain pairings shifted epitopes to avoid steric challenges posed by Omicron mutations. These findings support a model of anticipatory breadth with three key elements: (1) targeting of a functionally restricted epitope, (2) affinity maturation to establish an affinity buffer, and (3) variable chain pairing to generate paratope diversity. These elements each serve to compensate for a distinct consequence of viral mutagenesis, offering a mechanistic framework for anticipating viral evolution.

**Graphical Summary:** 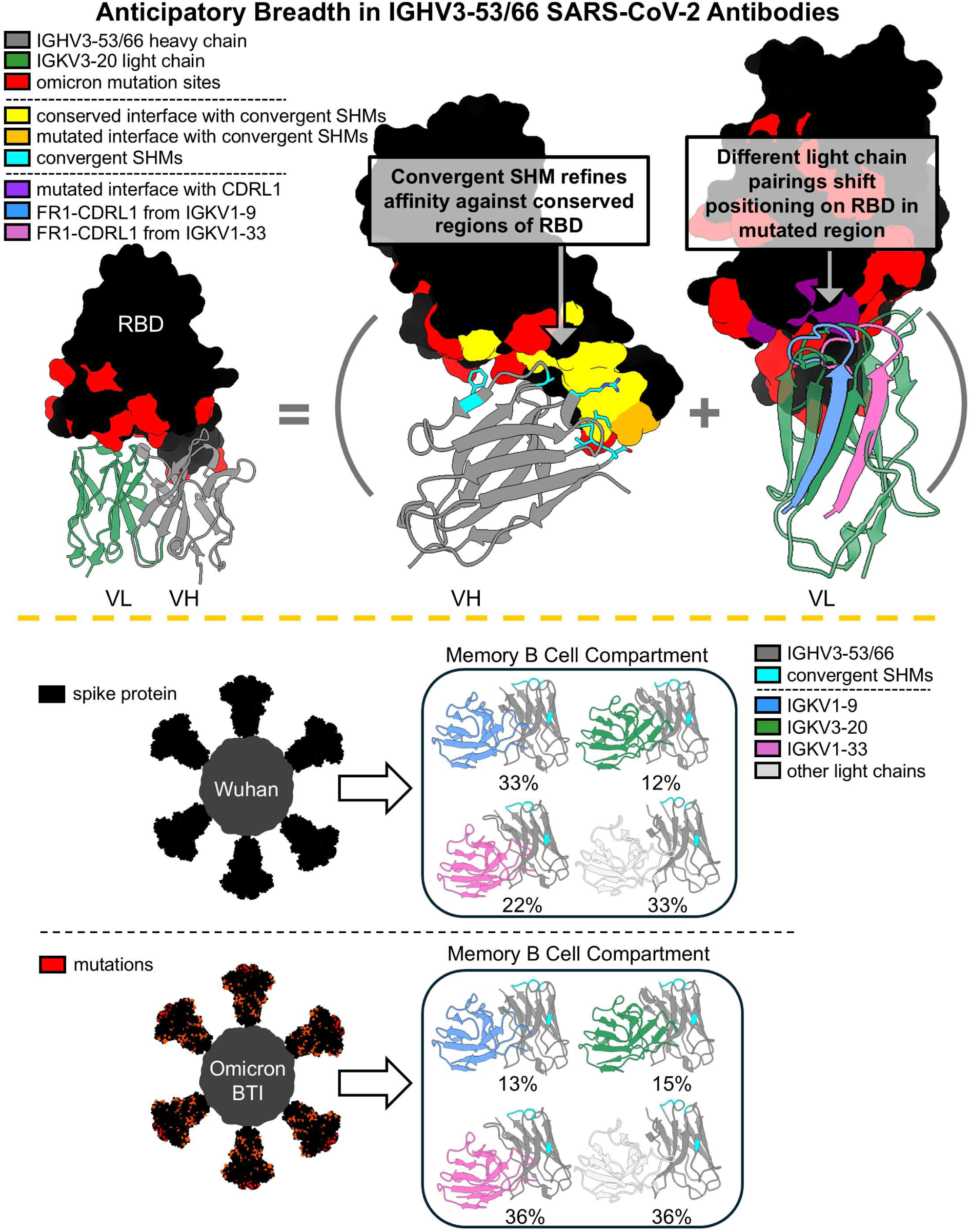

## Introduction

An optimal immune response would not only neutralize circulating pathogens but also anticipate future variants. In some cases, antibody responses can generate anticipatory breadth^1–8^, a phenomenon by which affinity maturation against ancestral strains produces antibodies capable of neutralizing future evolved strains. Understanding the mechanisms that generate anticipatory breadth is essential for designing vaccines that are resilient to viral evolution.

The human antibody response to SARS-CoV-2 offers a compelling model to study anticipatory breadth. A hallmark of this response is the strong elicitation of a public antibody family of potent neutralizers, defined by nearly identical heavy-chain genes IGHV3-53 and IGHV3-66^9–11^. IGHV3-53/66 antibodies bind a functionally restricted epitope on SARS-CoV-2 receptor binding domain (RBD) with extensive overlap with the binding site for ACE2^9,12^, the receptor required for viral entry into host cells. This binding is primarily mediated by germline-encoded residues^9,10^, allowing many IGHV3-53/66 antibodies to neutralize the ancestral Wuhan strain of SARS-CoV-2 with minimal refinement through somatic hypermutation (SHM)^13–15^.

Despite evolutionary constraints imposed by ACE2, the RBD has undergone extensive mutagenesis over time, including within the ACE2 binding region^16^. Nonetheless, several IGHV3-53/66 broadly neutralizing antibodies (bnAbs) have been identified, capable of neutralizing highly mutated Omicron strains^12,17–28^. These bnAbs typically exhibit higher levels of SHM than were observed early in the pandemic^24,25,28,29^. Additionally, several hallmark SHMs have been repeatedly identified across IGHV3-53/66 bnAbs, suggesting a convergence of breadth-conferring features^18,22,23,25,27,28,30–32^. While most reported IGHV3-53/66 broadly neutralizing antibodies (bnAbs) have been isolated from individuals who experienced breakthrough infections (BTI) — having been vaccinated against Wuhan SARS-CoV-2 and subsequently infected with Omicron — several IGHV3-53/66 bnAbs, as well as antibodies harboring hallmark SHMs, have also been isolated from individuals exposed only to the Wuhan strain^27,28,30,31^. The impact of variant exposure on breadth development, as well as the mechanisms by which breadth is achieved, remain unclear.

Anticipatory breadth is well established at the serum level for SARS-CoV-2; a standard two-dose mRNA vaccine series encoding the ancestral Wuhan spike, followed by a homologous booster administered 6+ months later, can elicit broadly neutralizing serum activity against Omicron-lineage variants^26,27,30–36^. To investigate mechanisms of anticipatory breadth development at the antibody level, we isolated IGHV3-53/66 antibodies from RBD-specific memory B cells of 9 infection-naïve individuals who had received two or three doses of the Wuhan mRNA vaccine. These antibodies exhibited broad neutralization against Omicron variants and displayed hallmark SHMs previously associated with neutralization breadth in BTI-derived antibodies^23,29,37,38^. Comparative analysis, leveraging available sequence, structural, and neutralization data, revealed both variant-exposure-associated and variant-exposure-independent features of breadth enhancement. Our findings suggest a model of anticipatory breadth relying on distinct mechanisms for combatting affinity loss from “receding” antigen interface, and steric challenge from “advancing” antigen interface, two distinct and general consequences of viral evolution.

## Results

### IGHV3-53/66 antibodies are a compelling case study for anticipatory breadth

We selected anti-SARS-CoV-2 IGHV3-53/66 antibodies as a model for studying anticipatory breadth based on 3 criteria: (1) targeting of a functionally constrained epitope, (2) frequent elicitation following infection or vaccination, and (3) demonstrated potential to develop into broad neutralizers. These criteria collectively constrain the potential for immune escape, ensure adequate sampling for analysis, and establish relevance to the overall neutralizing antibody response. While these criteria are generally established, we analyzed available sequence, structural, and functional data produced since 2020 for further evaluation.

We first profiled available structures to examine IGHV3-53/66 antibody epitopes. Of the 55 unique IGHV3-53/66 antibody:Wuhan RBD complexes in the Protein Data Bank (PDB), 51 engage a nearly identical epitope. This consistency in targeting contrasts with most other antibody families (Figure 1A), an observation corroborated by a large-scale epitope-profiling study using deep mutational scanning^29^ (Supplemental Figure 1). Notably, the 4 IGHV3-53/66:Wuhan RBD structures that exhibit alternative RBD binding have the largest CDR3s amongst all 55 structures, in line with a previous report linking long CDR3s to non-canonical binding for IGHV3-53/66 antibodies^38^. Overall, the high degree of epitopic convergence of IGHV3-53/66 antibodies supports reliable comparisons across the family.

**Figure 1.**
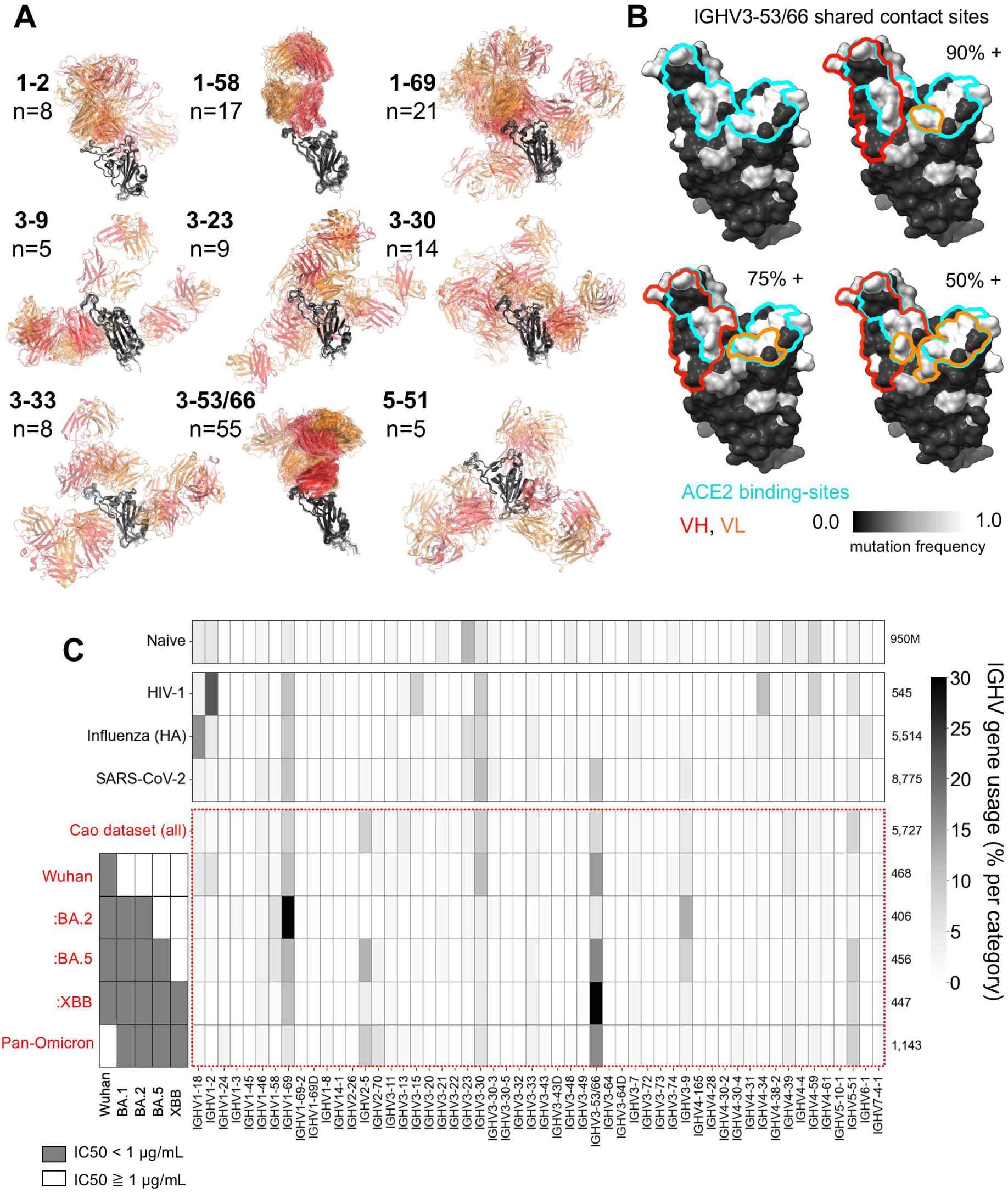
IGHV3-53/66 antibody epitope and usage frequencies. (A) Superposition (with respect to RBD) of antibody:Wuhan RBD complexes from the PDB, grouped by VH gene usage. Heavy chains are shown in red, light chains are shown in orange, and RBD is shown in black. (B) Shared epitope of 51 IGHV3-53/66 antibodies exhibiting “canonical” binding. Percentages indicate the fraction of antibodies that contact all RBD residues within highlighted VH and VL regions. RBD positions are colored based on sequence variability across variants of concern. (C) IGHV gene usage from published BCR datasets: Naive^63–65^, HIV-1 (CATNAP database^66^), Influenza (HA) (Wu Lab^67^), SARS-CoV-2 (CoV-Abdab^39^), Cao dataset (Cao^23,29,37,38^). Cao dataset is used to profile IGHV gene usage across neutralization categories spanning Omicron variants.

Within the canonical IGHV3-53/66 antibody epitope, heavy chain contacts are more consistent than light chain contacts. Across the 51 similar structures, heavy chains contact an average of 20.6 RBD residues, with 18 specific contacts shared by >90% of heavy chains. In contrast, light chains contact an average of 9.9 RBD residues, with only 4 specific contacts shared by >75% of light chains (Figure 1B). This profiling suggests that heavy chains provide a stable binding interface, while light chains may play a modular role. However, the regions targeted by both heavy and light chains are highly mutated across SARS-CoV-2 variants of concern (Figure 1B), potentially complicating relative contributions to neutralization breadth.

Next, we evaluated elicitation of IGHV3-53/66 antibodies by SARS-CoV-2, analyzing VH gene segment frequencies across several large datasets. Among all SARS-CoV-2-binding antibodies in CoV-AbDab^39^, the largest centralized database for SARS-CoV-2 antibodies, IGHV1-69, IGHV3-30, and IGHV3-53/66 are the most frequent VH genes utilized. Notably, IGHV3-53/66 appears to be specifically enriched among SARS-CoV-2 binders, whereas IGHV1-69 and IGHV3-30 gene segments show enrichment among binders to other viral pathogens such as HIV and influenza (Figure 1C).

We further examined IGHV3-53/66 gene frequencies across antibodies included in the largest published dataset of neutralization-tested, anti–SARS-CoV-2 monoclonal antibodies, here referred to as the Cao dataset^23,29,37,38^ (Figure 1C, Supplemental Table 1). This dataset contains nearly 6,000 antibodies from RBD-specific memory B cells of individuals who had received CoronaVac vaccination or experienced Omicron breakthrough infection. While IGHV3-53/66 antibodies accounts for only 7.4% of the full dataset, their frequency rises to 16.7% among antibodies that neutralize Wuhan through the Omicron BA.5 strain, and to 38.9% among those that neutralize Wuhan through Omicron XBB. Moreover, IGHV3-53/66 remains the most enriched VH family among bnAbs even when Wuhan-neutralizing capacity is not considered, at 16.4% (Figure 1C).

In summary, IGHV3-53/66 antibodies are frequently elicited following a specific viral exposure, exhibit consistent targeting of a restricted epitope, and make up the largest proportion of broad neutralizers targeting a common receptor binding domain. These features, in addition to prior observations of (1) robust anticipatory breadth in the context of Wuhan mRNA vaccination against SARS-CoV-2, and (2) individual IGHV3-53/66 bnAbs elicited in the absence of variant exposure, motivated our inquiry into the mechanism by which IGHV3-53/66 antibodies achieve anticipatory breadth against highly mutated SARS-CoV-2 variants.

### IGHV3-53/66 antibodies elicited by Wuhan mRNA vaccination neutralize Omicron variants and have known SHM hallmarks

To isolate antibodies of interest for neutralization profiling, we collected blood samples from 9 infection naïve, Wuhan-spike mRNA vaccinees 1 and 6 months after their second vaccine dose, and 10 days after their third, booster dose (Figure 2A, Supplemental Figure 2). Antibody sequences were cloned from sorted Wuhan RBD-specific memory B cells and binned into epitope groups by competition ELISA (Figure 2A). Binders overlapping with the ACE2 binding site (referred to as RBD-2 binders) were further evaluated for neutralization against Wuhan strain, as well as against a panel of Omicron variants, via pseudotype neutralization assay. Among 21 isolated RBD-2 binders, 11 were IGHV3-53/66 antibodies (Figure 2B). These IGHV3-53/66 antibodies generally had greater neutralization breadth than other RBD-2 binders, with 4 of 11 neutralizing all tested strains through XBB (Figure 2C).

**Figure 2.**
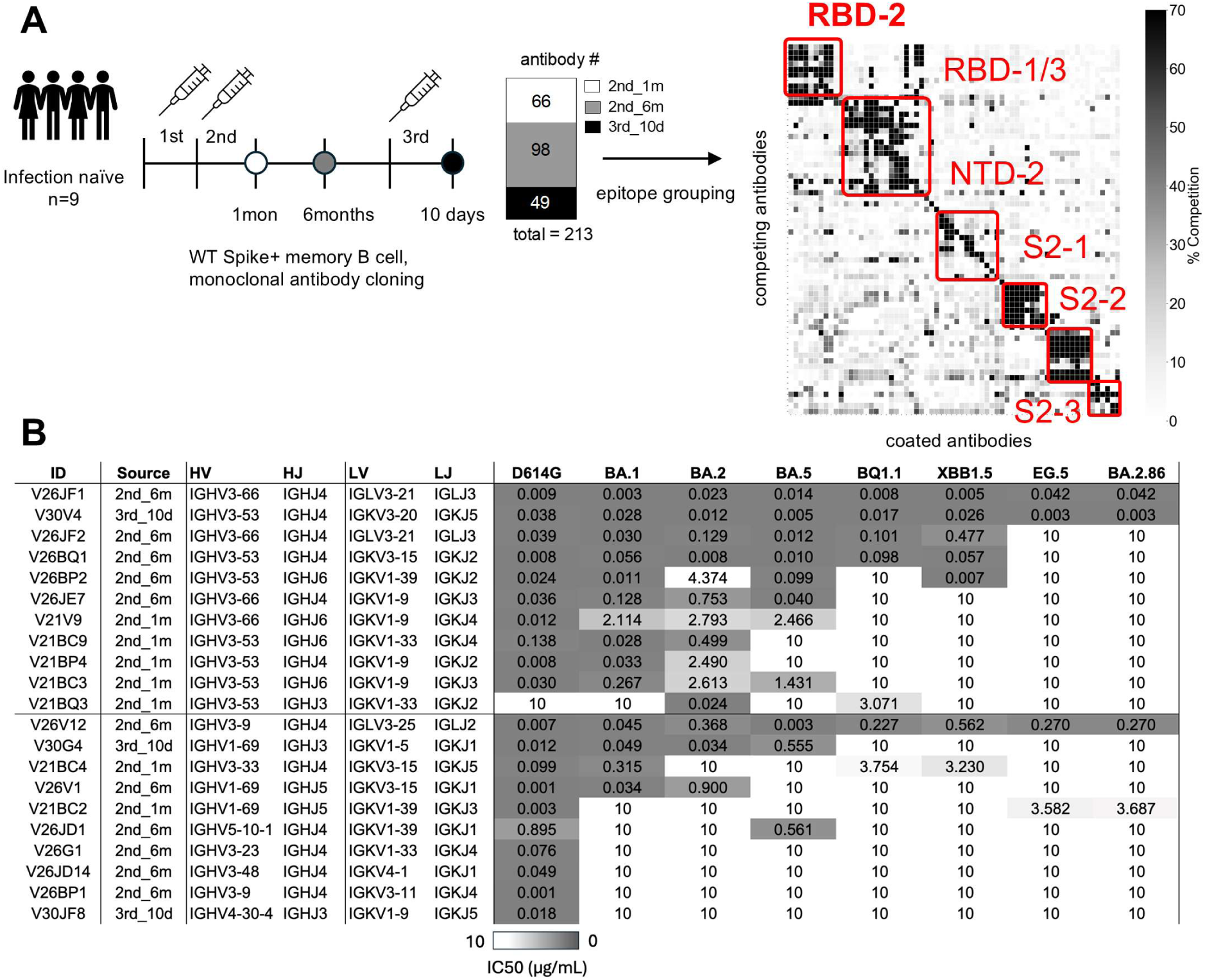
Isolation and neutralization testing of RBD2 antibodies from Wuhan spike mRNA vaccinees. (A) Study cohort and antibody isolation. Early pandemic SARS-CoV-2 mRNA vaccinees received 2 standard vaccine doses followed by a homologous booster dose. Antibodies were cloned from RBD-specific memory B cells collected across 3 timepoints. Expressed antibodies were grouped by epitope via competition ELISA. (B) RBD2 binders were further characterized for neutralization across a panel of SARS-CoV-2 variants by pseudotype neutralization assay.

To identify features associated with neutralization breadth, we assessed SHM, affinity to Wuhan RBD, and light chain usage. Six of the most described SHMs across IGHV3-53/66 antibodies— G27(26)E^18,27,30^, F28(27)I^28,40–42^, T29(28)I^27,30,40,43,32^, S36(31)R^28,30,32^ (CDRH1), and S58(53)P^28,30^,

Y66(58)F^30,38,42,32^ (CDRH2)—were present in our antibodies at varying frequencies (Figure 3, mutations are cited based on the IMGT numbering scheme^44^, with Kabat numerical identifiers in parentheses). Each of these mutations has been shown to increase affinity to Wuhan RBD, as well as to enhance neutralization breadth for individual tested IGHV3-53/66 antibodies by mutation reversion experiments^25,30,40–42^. Y66F was the most frequent mutation, present in 8/11 antibodies. G27E, the only mutation among the six to show a possible correlation with neutralization breadth in our dataset, was only present in the two most broadly neutralizing antibodies. F28I and S58P were less common than alternative mutations at their positions, F28V and S58A. A previous structural study supports enhanced complementarity of the CDR1-RBD interface by F28I that also applies to F28V^42^. Although the impact of the S58A mutation is less clear, its positioning and frequency suggests that removal of serine at position 58 may be favorable.

**Figure 3.**
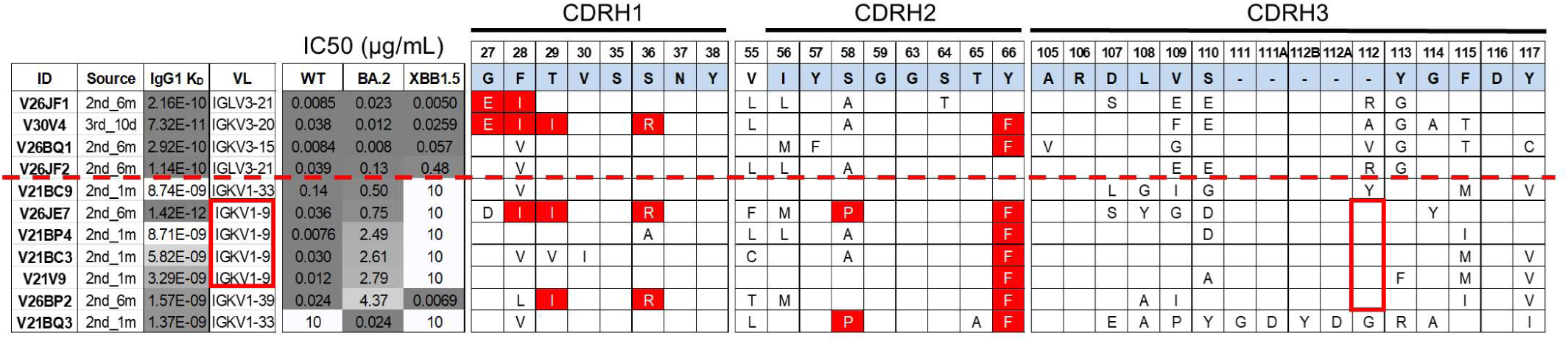
Features of IGHV3-53/66 antibodies isolated from Wuhan spike mRNA vaccinees. 11 IGHV3-53/66 monoclonal antibodies (mAbs) were profiled for affinity for Wuhan RBD, light chain pairing, neutralization breadth, and VH segment somatic hypermutation. Antibodies above the dashed red line neutralize all tested strains of SARS-CoV-2 through XBB. Red brackets highlight features with apparent enrichment among poorer neutralizers. Sequence numbers are based on the IMGT numbering scheme^44^. Germline amino acids across CDRH1 and CDRH2 and common amino acids across CDRH3 are noted in light blue on the top row above mAb sequences. Blank boxes across mAb sequences represent amino acid agreement with germline/common residues. Amino acids in red boxes are previously identified, “hallmark” SHMs shown to have positive impacts on neutralization breadth and affinity to Wuhan RBD.

Biolayer interferometry revealed a general correlation between affinity for Wuhan RBD and neutralization breadth. Four of the five highest-affinity antibodies neutralized all strains through XBB (Figure 3). A notable exception was V26JE7, which despite being the highest-affinity binder, exhibited comparatively poor neutralization values and did not neutralize the XBB strain. This antibody has a IGKV1-9 light chain and a CDRH3 length of 11 amino acids—both features associated with limited breadth among our IGHV3-53/66 antibodies (Figure 3).

Together, these results demonstrate that infection-naïve individuals receiving Wuhan mRNA vaccines can generate IGHV3-53/66 antibodies with broad neutralization capacity and SHM hallmarks previously associated with neutralization breadth.

### Convergent SHM refines germline-encoded RBD contacts at conserved sites

One of the broadest neutralizers in our cohort, V30V4, contained all 6 previously described hallmark SHMs (with S58A rather than S58P), and had striking sequence similarity (including the same VL gene pairing) to Omi-3, a previously reported bnAb isolated from a BTI sample^17^ (Figure 4A). To investigate how hallmark SHMs relate to epitopic changes between Wuhan and Omicron RBDs, we determined the structure of V30V4 in complex with Wuhan RBD by Cryo-EM (extra methods description included in supplementary information) and compared it to an available structure of Omi-3 in complex with BA.1 Omicron RBD (PDB ID: 7ZF3).

**Figure 4.**
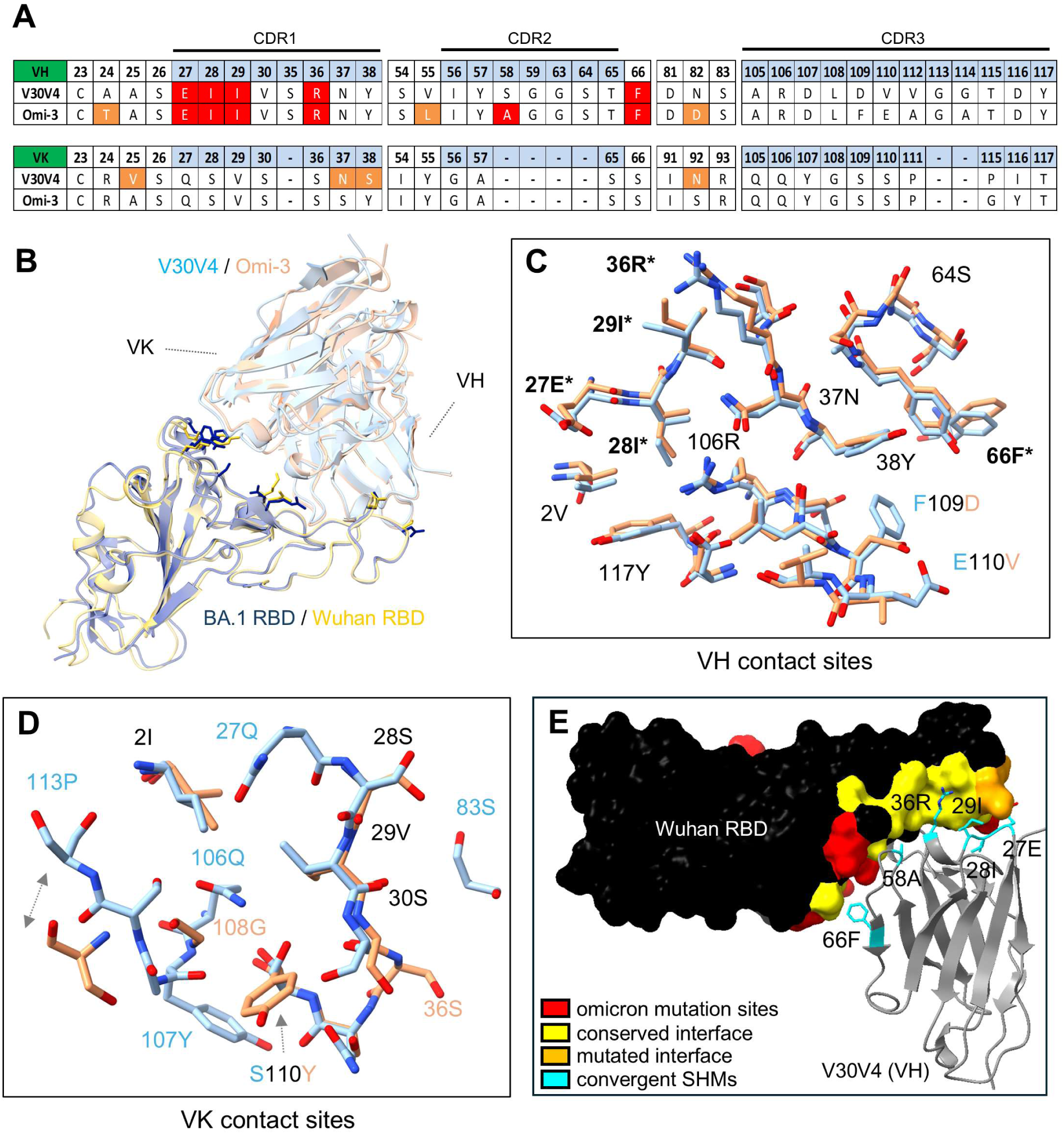
Comparison of V30V4 to Omi-3 and profiling of V30V4 “hallmark” SHM contacts. (A) Sequence comparison of 2 similar antibodies: V30V4, isolated from our cohort of Wuhan mRNA vacinees, and Omi-3, isolated from an Omicron BTI sample. Sequence numbers are based on the IMGT numbering scheme^44^. Hallmark SHMs for IGHV3-53/66 antibodies are highlighted in red. Additional SHMs are highlighted in orange. (B) Structures of V30V4 and Omi-3 superimposed with respect to RBD. Residues at mutation sites distinguishing Wuhan and BA.1 RBD structures and involved in antibody interfaces are visualized. (C-D) Positional comparison between V30V4 and Omi-3 contact residues (superposition with respect to RBD) are visualized for (C) VH, and for (D) VL. (E) The structure of V30V4 VH in complex with Wuhan RBD is presented, highlighting the contact interface between hallmark SHMs and Wuhan RBD. Non-interfacing RBD positions that mutate in Omicron strains are shown in red.

Several Omicron mutations alter the antibody:RBD interface by either adding or removing contact area. K417N and Y505H are known escape mutations that reduce affinity of the RBD for IGHV3-53/66 antibodies through “receding” interface^20,25,27,43^. Conversely, N501Y results in “advancing” interface, modifying the steric relationship between RBD and IGHV3-53/66 paired light chains. Accordingly, Omi-3 has less contact with BA.1 RBD at positions 417 and 505 than the V30V4:Wuhan RBD complex, and more contact at position 501 (Supplemental Figure 3). Despite these differences, docking of the Fab on RBD is very similar between V30V4 and Omi-3. At the chain level, VH positioning is nearly identical between V30V4 and Omi-3 (Figure 4B), with a root-mean-squared distance (RMSD) of 0.819 angstroms (superposition with respect to RBD). This similarity is also reflected in the nearly identical positioning and orientation of their VH contact residues (Figure 4C). VL positioning is less similar (Figure 4B), with an RMSD of 1.714 angstroms, falling to 1.283 angstroms when only considering residues of structurally conserved beta sheets. Correspondingly, VL contact residues also overlap less (Figure 4D), likely reflecting more conformational compensation for RBD mutations on the VL side than on the VH side. Overall, these structures show that minimal conformational compensation, especially by VH, may be required for effective engagement of Omicron RBD by some Wuhan-elicited IGHV3-53/66 antibodies.

We also profiled contact sites between RBD and hallmark SHMs across V30V4. Hallmark SHMs mostly associate with conserved regions of RBD, with 10/12 contact residues conserved across Wuhan and variant strains (Figure 4E). Comparison with an available structure of Wuhan RBD complexed with an IGHV3-53/66 antibody with no SHM and the same light chain pairing as V30V4 (PDB ID: 7D0C) further revealed that 4 out of 6 hallmark SHMs refine exclusively pre-existing germline-encoded contacts (Supplemental Figure 4). Only 2 mutations, G27E and S36R, establish new contacts with RBD, including that between G27E and the only 2 non-conserved residues engaged by any of the hallmark SHMs (Supplemental Figure 4B). These observations suggest that the high degree of germline complementarity between IGHV3-53/66 antibodies and Wuhan RBD may facilitate convergent SHM, and that refinement of affinity against conserved regions of RBD by SHM may help preserve epitope targeting and binding orientation despite interface-altering mutations in the RBD.

### SHM associated with neutralization breadth arises independently of variant exposure

Hallmark SHMs, frequent across our cohort, target conserved regions of RBD and appear consistently across reported BTI-derived IGHV3-53/66 antibodies^21,22,25^. However, whether exposure to drifted variants influences overall SHM or results in selection of key breadth-enhancing mutations remains unclear. To evaluate the relationship between variant exposure and neutralization breadth development for IGHV3-53/66 antibodies, we analyzed the Cao dataset, which includes full antibody sequences and neutralization profiles through the XBB variant for 179 IGHV3-53/66 antibodies (Supplemental Table 1). To isolate only high-confidence binders to the canonical IGHV3-53/66 epitope, we filtered for antibodies with CDRH3s ≤15 amino acids, as long CDRH3s have been previously associated with alternative epitope targeting^45^. After filtering, approximately half of the remaining 156 IGHV3-53/66 antibodies are derived from Omicron breakthrough infection (BTI) samples, while the rest come from individuals exposed only to the ancestral Wuhan strain through infection or CoronaVac vaccination.

We grouped antibodies in two ways (Figure 5A): (1) by breadth, comparing bnAbs (neutralizing Wuhan through XBB) to narrow neutralizers (Wuhan only), and (2) by exposure, comparing BTI (Omicron-exposed) samples to Wuhan-only exposed samples. Initial comparison of antibodies by exposure group revealed higher frequencies of broad neutralizers in the BTI cohort: roughly 57% and 16.5% of BTI-derived IGHV3-53/66 antibodies neutralize all strains through BA.5 and XBB, respectively, compared to 20.8% and 6.5% in the Wuhan-exposed group (Supplemental Figure 5A). However, BTI antibodies also have higher average VH-region SHM (7.9 mutations, Supplemental Figure 5B) than the Wuhan-only group (5.0 mutations, Supplemental Figure 5C), suggesting that antibodies from the BTI group were subjected to greater affinity maturation.

**Figure 5.**
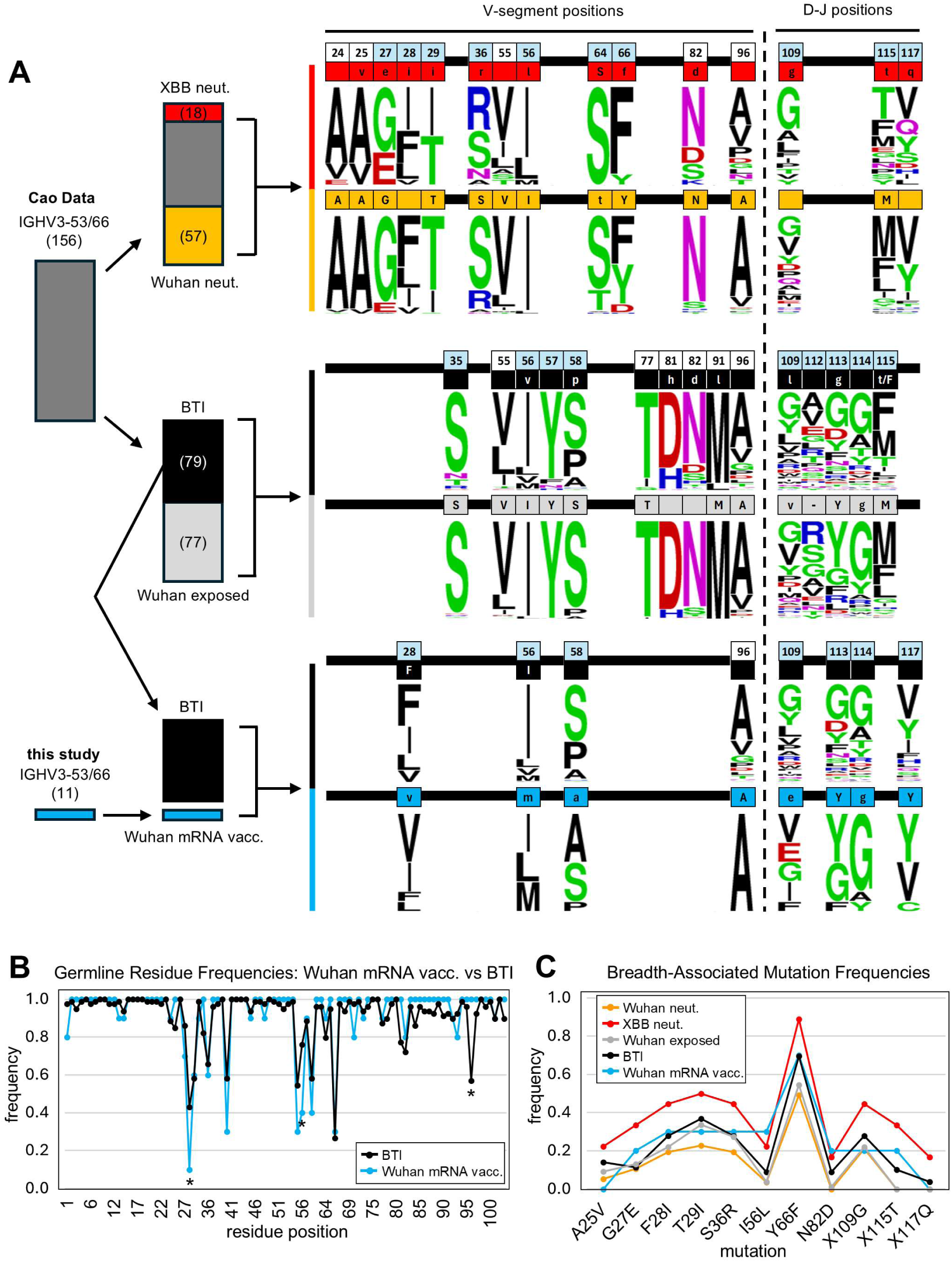
Sequence analysis of IGHV3-53/66 antibodies by neutralization breadth and exposure categories. (A) Schematic detailing analysis of sequenced IGHV3-53/66 antibodies from the Coa dataset^29,37^ and subsequent comparison to Wuhan-mRNA derived IGHV3-53/66 antibodies from this study. Antibodies with CDRH3 length >15 were excluded from analysis. IGHV3-53/66 antibodies from the Cao dataset were divided into groups based on neutralization breadth or exposure. For each group, significantly enriched positional amino acids were determined by Fisher’s exact test (p <0.05). Significant positions were then used to compare BTI sequences from the Cao dataset to Wuhan-mRNA derived IGHV3-53/66 antibodies from this study. Enriched positional amino acids are noted in color-coded boxes. Overall amino acid diversity at each position for each group is illustrated by logo plot. CDR positions are noted in light blue boxes. Sequence numbers are based on the IMGT numbering scheme^44^. Germline residues are denoted by capital letters, and mutations are denoted by lower-case letters. (B) Line graph illustrates germline residue frequency across IGHV3-53/66 antibodies from Cao BTI (black) and this study (blue). Asterisks denote significant differences (Fisher’s exact test, p < 0.05). (C) Line graph illustrates frequencies of specific mutations enriched in broad neutralizers (XBB neut. category) across noted antibody groups. No significant differences are observed between Cao BTI antibodies and antibodies from this study across these mutations.

Turning to sequence analysis, we identified 12 VH positions that exhibit statistically significant differences in specific amino acid frequencies between broad and narrow neutralizers, and 10 differential positions by exposure group (Fisher’s exact test, p<0.05). Four positions—55, 56, 82, and 96—overlap between the two comparisons. Notably, 5 of the 6 previously described hallmark mutations—G27E, F28I, T29I, S36R, and Y66F—are enriched in broad neutralizers but not significantly different between BTI and Wuhan-exposed groups. Given lower levels of SHM for the Cao Wuhan-exposed group compared to the BTI group, lack of variation of hallmark mutations between these groups is strong evidence for highly convergent SHM independent of variant exposure.

Due to the diversity of D and J gene usage, and of CDRH3 structure, our power to detect statistically significant sequence differences in the DH-JH region is limited. However, one residue,115T, has the strongest statistical association with broad neutralizers of any residue across the entire VH sequence, and is also enriched in the BTI group (Figure 5A). This residue is also present in 2 of our XBB-neutralizing Wuhan mRNA-derived antibodies (Figure 3); 115T, while not directly involved in the interface with RBD, replaces other larger, hydrophobic residues at the base of CDRH3 that may restrict CDRH3 positioning (Figure 5A, Supplemental Figure 6A). Its appearance in our small cohort, despite extra variability in the CDRH3 region, suggests a strong selection for this residue, independent of variant exposure.

We also compared our IGHV3-53/66 antibodies from Wuhan mRNA vaccinees to BTI-derived sequences from the Cao dataset. Of the 18 V-gene positions with statistically significant differences by neutralization and/or exposure groups across Cao data, only 4 differed significantly between our sequences and Cao BTI sequences (Figure 5A). Three of the four were associated with neutralization breadth (positions 28, 56, and 96), and one with exposure (position 58). For positions 28, 56, and 58, BTI sequences were enriched for germline residues more common among narrow neutralizers and Wuhan-exposed samples, reflecting mildly enhanced breadth among our Wuhan mRNA-derived antibodies.

At position 96—among the most frequently mutated sites in BTI sequences—our Wuhan mRNA-derived antibodies had a mutation frequency of 0 (Figure 5B). This site resides at the base of framework region 3 (FRH3), far from the antigen interface, and is unlikely to impact binding (Supplemental Figure 6B). Even within the Wuhan-exposed subset of the Cao dataset, this position showed frequent SHM, suggesting that mutation at position 96 may be cohort-specific. While not statistically significant, SHM was also absent from positions 83–91 in our Wuhan mRNA-derived antibodies compared to mild levels in BTI antibodies (Figure 5B). This region corresponds to a beta sheet on the back of FRH3, unlikely to contribute to antigen interaction (Supplemental Figure 6B). Finally, positions 70 and 93—both on the solvent-facing side of FRH3—showed higher mutation frequency in our Wuhan mRNA-derived antibodies compared to BTI antibodies (Figure 5B), despite having no known role in RBD recognition (Supplemental Figure 6B). Across all specific breadth-associated SHMs defined by comparison of broad and narrow neutralizers in the Cao dataset, there were no significant differences between Cao BTI antibodies and our Wuhan mRNA-derived antibodies (Figure 5C).

These comparisons illustrate that SHM associated with neutralization breadth occurs independently of variant exposure, supporting a model in which neutralization breadth develops primarily through affinity maturation against conserved features of RBD.

### Correlation with light chain pairing confounds the relationship between CDRH3 length and breadth

In our Wuhan mRNA-derived IGHV3-53/66 antibodies, we observed two features associated with narrow neutralization: a CDRH3 length of 11 amino acids and pairing with IGKV1-9 light chain (Figure 3). To explore these features further, we analyzed their distribution across exposure and neutralization groups in the Cao dataset.

The frequency of IGHV3-53/66 antibodies with 11-residue CDRH3s decreases with increasing variant exposure and neutralization breadth. This decrease coincides with a rise in antibodies with 12-residue CDRH3s (Supplemental Figure 7A). A similar trend was observed for light chain usage; IGKV1-9 pairing is enriched among Wuhan-only exposed and narrowly-neutralizing antibodies, while IGKV1-33 pairing is enriched among Omicron-exposed and broadly neutralizing antibodies (Figure 6A). However, because IGKV1-9 usage is also positively correlated with 11-residue CDRH3s (Supplemental Figure 7B), the relative contribution of each variable to breadth is difficult to assess. The proportion of IGKV1-9 paired antibodies with 11-residue CDRH3s is similar between exposure groups (Supplemental Figure 7C), suggesting that features beyond CDRH3 length may drive the apparent negative selection of IGKV1-9 paired antibodies by variant exposure.

**Figure 6.**
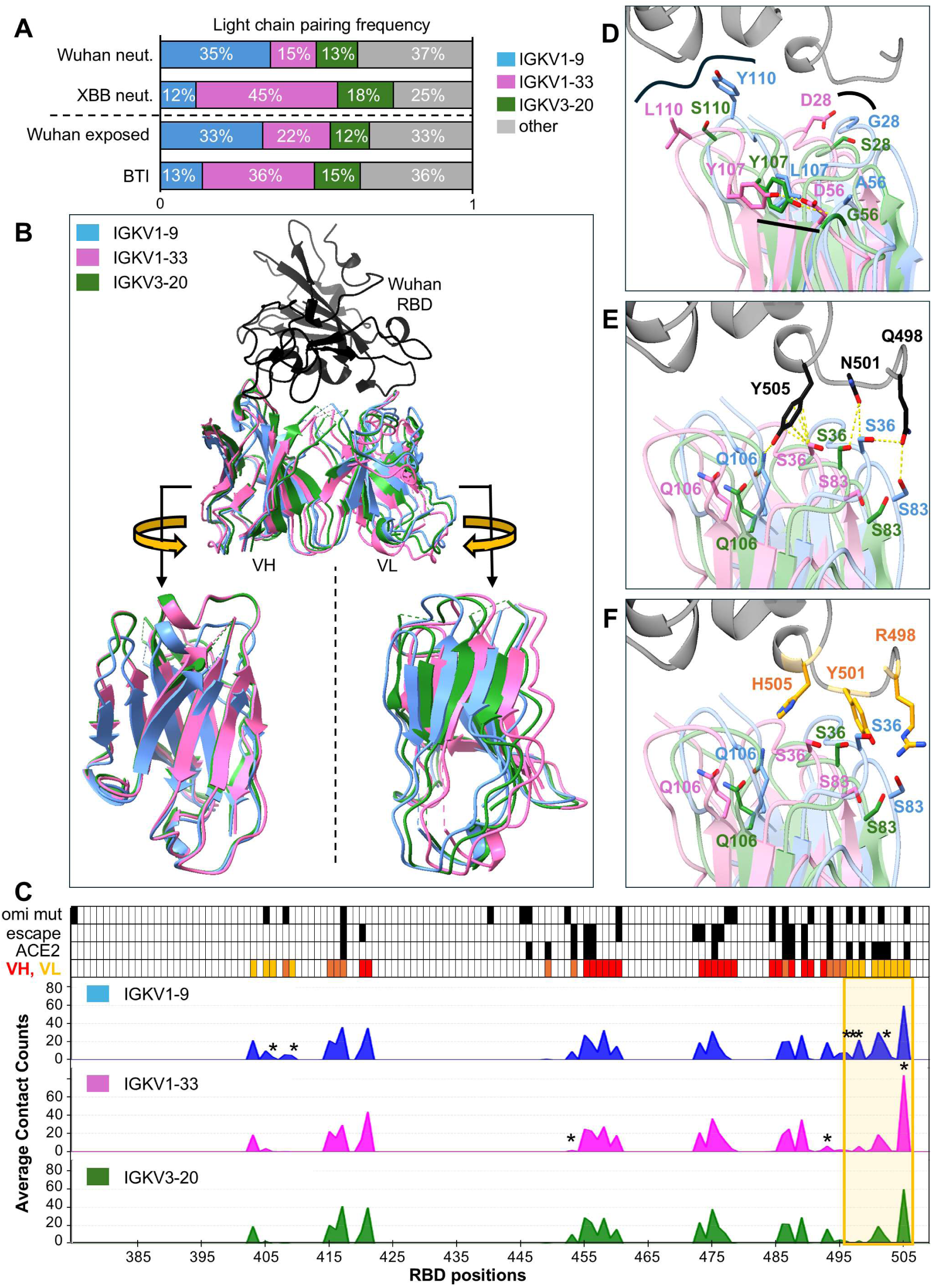
Omicron exposure selects for favorable light chain pairing for IGHV3-53/66 antibodies. (A) Light chain gene frequencies for IGHV3-53/66 antibodies from the full Cao dataset^23,29,37,38^ across neutralization and exposure categories. (B) Superposition (with respect to RBD) of average backbone structures of light-chain-defined groups of available IGHV3-53/66 antibody:Wuhan RBD PDBs. Structures were generated by calculating the average coordinate for each common backbone atom across available PDB structures of IGHV3-53/66 antibodies paired with common light chains and complexed with Wuhan RBD. Rotated views of heavy and light chains highlight spatial shifts between light chain defined groups. (C) Plotted averages of atomic contacts with RBD positions by light-chain-defined groups of IGHV3-53/66 antibodies from available PDBs. Asterisks denote significant differences (p <0.05). Top rows illustrate positions of Omicron mutations (“omi mut”), escape mutations (“escape”, based on DMS data^29^), ACE2 binding (“ACE 2”), and contact with VH and/or VL (“VH, VL”). Gold box highlights most differential region for RBD contact. (D-F) Representative structures of commonly paired IGHV3-53/66 antibody light chains in complex with RBD, superimposed with respect to RBD (PDB IDs: 7Q0A (pink), 7B3O (blue), 6XC7 (green)). (D) Germline differences between light chains contribute to steric shifts. (E) Steric shifts contribute to variable RBD contacts with shared germline light chain residues. (F) Pan-Omicron mutations modify steric and interfacial relationships with different paired light chains for IGHV3-53/66 antibodies. Omicron RBD (BA.1) from Omi-3:RBD structure (PDB ID: 7ZF3).

### Light chain pairing generates anticipatory breadth through paratope diversification

To evaluate relationships between light chain pairing, variant exposure, and neutralization breadth, we first calculated frequencies of the 3 most common light chain genes paired with IGHV3-53/66 antibodies—IGKV1-9, IGKV1-33, and IGKV3-20—in the Cao dataset across exposure and neutralization groups (Figure 6A). IGKV1-9 is the most frequent light chain pairing for IGHV3-53/66 antibodies amongst Wuhan-exposed and narrow neutralizers, and the least frequent pairing among BTI-derived and broad neutralizers. Conversely, IGKV1-33 is the most frequent light chain pairing for IGHV3-53/66 antibodies amongst BTI-derived and broad neutralizers, and the least frequent pairing among Wuhan-exposed and narrow neutralizers.

IGKV3-20 frequencies remain stable (∼10-20%) across exposure and breadth categories. These trends point to a strong selection for IGKV1-33 light chains by Omicron exposure, suggesting that IGKV1-33 light chain pairing may be advantageous for engaging Omicron RBD.

To investigate the structural consequences of light chain pairing, we compiled PDB structures of IGHV3-53/66 antibodies paired with IGKV1-9, IGKV1-33, and IGKV3-20 light chains in complex with Wuhan RBD. Only antibodies with CDRH3 lengths ≤13 were included to minimize structural variability (IGKV1-9, n = 14, IGKV1-33, n = 7, IGKV3-20, n = 13). After superimposing all structures with respect to RBD, we noted that light chain groups formed distinct, overlapping clusters (Supplemental Figure 8A), suggesting that light chain pairing may influence overall antibody docking.

To quantify spatial differences, we calculated the average structure of each light-chain-defined group. Average structures revealed a clear gradient in light chain positioning, with IGKV1-9 most peripheral on RBD, IGKV1-33 most central, and IGKV3-20 occupying the middle of the gradient (Figure 6B). RMSDs between average structures reflected this gradient, with IGKV1-9 and IGKV1-33 separated by ∼3 Å, and IGKV3-20 nearly equidistant from both (Supplemental Figure 8B). While heavy chain positioning was similar between average IGKV1-33-paired and IGKV3-20-paired structures, IGKV1-9 pairing induced a notable downward shift in CDRH1 and CDRH2 away from RBD (Figure 6B). These spatial differences led us to examine atomic-level contacts between light-chain-defined groups.

Wuhan RBD contact profiles for light-chain-defined groups revealed modest differences in RBD contacts across most of the RBD (Figure 6C). Contacts between RBD and heavy chains of light-chain-defined groups are particularly consistent. However, there are large differences in light chain contacts between light-chain-defined groups and RBD positions 496-505 (Figure 6C). This stretch of RBD is of particular interest, as it encompasses sites for 3 pan-Omicron mutations: Q498R, N501Y, and Y505H.

Close examination of representative structures of IGHV3-53/66 antibodies paired with IGKV1-9 (PDB ID: 7B3O), IGKV1-33 (PDB ID: 7Q0A), and IGKV3-20 (PDB ID: 6XC7) light chains highlight germline-encoded features that explain much of their distinct spatial positioning (Figure 6D). G28 in IGKV1-9 facilitates close CDRL1–RBD proximity, while D28 in IGKV1-33 increases separation. In CDRL3, Y110 in IGKV1-9 enables unique RBD contacts, although these contacts vary between antibodies. Intra-chain constraint on CDRL3, imposed by Y107 interaction with CDRL2 in IGKV1-33 and IGKV3-20, is broken by L107 in IGKV1-9. This constraint is modified by the broader separation between Y107 and D56 in IGKV1-33, compared to Y107’s interaction with G56 in IGKV3-20.

These structural differences lead to distinct RBD interactions. Despite all 3 light chains sharing germline residues at positions 36, 83, and 106, IGKV1-9 forms unique contacts with Q498 (via S36 and S83) and Y505 (via Q106) of RBD. In contrast, IGKV1-33 aligns for increased S36 contact at Y505 (Figure 6E). Across analyzed PDBs, IGKV1-9 forms the most contact with Wuhan RBD, with an average buried surface interface (BSA) of 457.3 Å^2^, compared to 316.4 Å^2^ for IGKV3-20, and 282.1 Å^2^ for IGKV1-33 (Figure 6C, Supplemental Figure 9, Supplemental Table 2). However, its positional advantage in the context of Wuhan RBD becomes a liability with the introduction of Omicron mutations. N501Y introduces a steric clash with CDRL1 of IGKV1-9, while it improves proximity with IGKV1-33 and IGKV3-20 (Figure 6F). Furthermore, CDRL1 in IGKV1-33 is best positioned to maintain contact with Y505H, a known escape mutation.

In summary, light chain pairing introduces paratope diversity into the IGHV3-53/66 antibody repertoire. This diversity allows some pairings, such as IGKV1-33, to better accommodate epitope remodeling caused by viral evolution, while others, such as IGKV1-9, may optimize interactions with the original antigen but face steric constraints upon drift. These findings support a model in which light chain–dependent diversification enables IGHV3-53/66 antibodies to balance ancestral affinity with future adaptability.

## Discussion

SARS-CoV-2 presented a unique opportunity to investigate how an initial immune response to a viral challenge can generate breadth against future variants. The inherent complementarity between human IGHV3-53/66 antibodies and the mutationally restricted ACE2 binding site on Wuhan RBD resulted in strong elicitation against the ancestral strain and development of broad neutralization against drifted strains. This phenomenon, amplified by a global immunization effort and extensive structural and functional characterization of related monoclonal antibodies, created a fortuitous model system for dissecting mechanisms that can give rise to anticipatory neutralization breadth to future evolved viral variants.

We identify two mechanisms by which IGHV3-53/66 antibodies develop neutralization breadth in the absence of variant exposure, (1) affinity maturation via SHM and (2) paratope diversification through variable light chain pairing. SHM refines contact with conserved epitopic regions within the ACE2 binding site, producing affinity gains that carry over to drifted variants. Meanwhile, variable light chain pairing, an inherent mechanism for diversifying the naïve antibody repertoire, modulates antibody-antigen docking to accommodate epitope remodeling. While the generalizability of this model remains to be evaluated, these findings support a framework for anticipatory breadth in which distinct mechanisms offset specific consequences of viral evolution.

Two types of viral mutations can be especially disruptive to antibody-antigen interfaces: those that (1) “recede” the epitope surface, reducing contact area, or (2) “advance” it, introducing steric clashes. For example, the pan-Omicron mutation K417N removes a bulky side chain and several key contact points for IGHV3-53/66 antibodies, decreasing affinity. In contrast, N501Y introduces a bulkier residue, clashing with the preferred docking site of IGKV1-9-paired antibodies. In our model, SHM provides an affinity buffer that offsets contact loss from receding mutations, while paratope diversity from variable chain pairing enables a subset of antibodies to avoid steric challenges from advancing mutations. Together, these mechanisms generate resilience to two major implications of viral drift.

Previous reports have highlighted additional mechanisms that may contribute to anticipatory breadth development in some cases, such as diversification through “bystander” mutagenesis observed in IGHV1-58 antibodies against SARS-CoV-2^46^. Independent of our model’s generalizability, the IGHV3-53/66 antibody case study illustrates an important principle for vaccine design. Comparative analysis of IGHV3-53/66 antibodies from Wuhan mRNA vaccinees and BTI samples revealed highly convergent SHM patterns that emerged independent of variant exposure. This is in line with prior evidence invoking the phenomenon of “original antigenic sin”^47^, suggesting that BTI in SARS-CoV-2 primarily results in recall and expansion of pre-existing, cross-reactive memory B cells, with minimal *de novo* response from elicited naïve B cells^32^. One would expect such selective activation to enrich for clear, differentiable features necessary for cross-reactivity. That frequencies of somatic mutations across IGHV3-53/66 antibodies remain largely unchanged following BTI highlights a level of convergence of the affinity maturation process that, despite having been noted previously^21,22,25^, may still have been under-appreciated.

This convergence may be due, in part, to the unusually high germline-encoded complementarity between IGHV3-53/66 antibodies and Wuhan RBD, favoring refinement of existing contacts over generation of new interface. Additionally, complementarity may play a role in the robust production of IGHV3-53/66 memory B cells following Wuhan exposure, seeding a diverse pool of light chain pairings that can be selectively expanded upon future antigenic challenge. In total, anticipatory breadth development in IGHV3-53/66 antibodies may highlight the importance of developing ideal immunogens with high levels of complementarity to intended germline antibody targets, potentially leading to diverse memory deposition, as well as predictable pathways of affinity maturation.

## Acknowledgments

We thank the volunteers who donated blood for our study. This study was supported by NIH grants AI139538, AI169619, AI170715, AI170580, and AI165072 to D.R.W. J.D. and Z.A. are supported by the National Institute of Arthritis and Musculoskeletal and Skin Diseases of the NIH under award number T32AR007530. J.W.C. is supported by NSF GRFP (DGE 2140743), NIH F31AI186232, and FujiFilm Fellowship. We are grateful to Stephen C. Harrison for providing comments on the manuscript.

## Author Contributions

D.R.W., J.D., D.K.Y., S.K. designed the study and analyzed data. J.D., D.K.Y., S.K., Z.A., J.C. conducted experiments and analyzed data. G.K., Y.W., and B.C. performed and analyzed cryo-electron microscopy data. Z.A. and S.S. performed computational and data analysis. J.D. wrote the manuscript with input from other authors. D.R.W. supervised the research and revised the paper.

## Materials and Methods

### Human participants

All procedures involving human samples were approved by the Massachusetts General Brigham (MGB) Institutional Review Board (Protocol #:2020P000837). Informed consent was collected from participants in this study. Blood samples were collected from COVID-19 infection naive individuals vaccinated with the COVID-19 mRNA vaccine (n=9). A longitudinal study was conducted where blood was collected one month after the COVID-19 booster (second dose), six months after the COVID-19 booster (second dose), and 10 days after the second booster (third dose). Vaccinated participants in this study self-reported to have no history of COVID-19 diagnoses.

### Plasma and peripheral blood mononuclear cell (PBMC) isolation

Blood collected from participants by our clinical team was collected in EDTA tubes. Blood was centrifuged at 300 xg for 10 minutes. Plasma was collected from the top layer and further centrifuged at 1000 xg for 10 minutes for the removal of debris. Plasma was divided into aliquots and stored in -80°C.

PBMCs were collected from the lower layer of the initial centrifuge step. Ficoll-Paque (Cytiva) was added to the lower layer for PBMC separation at 2200 rpm for 20 minutes. PBMCs were washed with PBS and resuspended in FBS containing 10% dimethyl sulfoxide (DMSO).

Samples were aliquoted and stored at -80°C and liquid nitrogen.

### Plasma nucleocapsid (N), SARS-CoV-2 spike subdomain protein reactivity ELISA

As previously described^48^, to test the infection status of the native SARS-CoV-2 virus, quantification of plasma IgG against SARS-CoV-2 nucleocapsid (N) protein (Genscript) was performed. In brief, 50 ng/well of the N protein, of the S protein, or 2 μg of the RBD protein in 30 μL of PBS were used to coat 96-well MaxiSorp ELISA plates (ThermoFisher Scientific). Plates were incubated overnight at 4°C. Coating solution was discarded after incubation and blocking buffer (100 μL of 3% bovine serum albumin, BSA) was applied to the plates for 2 hours. During incubation, plasma from SARS-CoV-2 vaccinated patients were inactivated. Plasma samples were incubated with equal volume of 2% Triton in PBS and incubated at room temperature for 20 minutes. Inactivated plasma was serial diluted from 1:200 to 1:1600. The mAb CR3022 was used as the standard for anti-S and anti-RBD. After incubation, the blocking buffer was discarded and the ELISA plates were washed once in washing buffer (PBS containing 0.05% Tween 20). Plasma dilutions were added to the ELISA plates in duplicates. 8 negative control sera (ProMedDx) from pre-pandemic period were purchased and included in the experiment. Plates were incubated at 4°C overnight.

After overnight incubation, the ELISA plates were washed three times in washing buffer before incubation for 90 minutes in 1:1000 dilution of alkaline phosphatase-conjugated anti-human IgG (Southern Biotech) in PBS containing 1% BSA and 0.05% Tween 20. After incubation, ELISA plates were washed three times with washing buffer then incubated in alkaline phosphatase substrate solution (Sigma-Aldrich) for two hours.

The ELISA plates were read at 405 nm by a microplate reader (Biotek Synergy H1) and GraphPad Prism v10 was used to generate standard curves for each plate by non-linear regression.

### Flow Cytometry and Single-cell sorting

PBMCs stored at -80°C were thawed at 37°C and transferred to warm RPMI 1640 media (Gibco) supplemented with 10% FBS (Cytiva). B cells were enriched by positive selection of B cells with anti-CD19 beads (Milytenyi). As previously described^49^, spike positive memory B cells were sorted (BD FACSAria Fusion) into 96-well plates. In brief, enriched B cells were stained with SARS-CoV-2 containing a Flag tag (Genscript). Following primary incubation, the enriched B cells were stained with allophycocyanin (APC)-conjugated anti-Flag and phycoerythrin (PE)-conjugated anti-Flag for double positive sorting. DAPI- IgM- IgD- IgG+ CD27+ Spike+ B cells were single cell sorted into 96-well PCR plates containing lysis buffer (4 μL of 0.5x PBS with 10 mM dithiothreitol and 4U of RNAseOUT). Plates were stored at -80°C.

### Antibody sequencing, cloning, and expression

As previously described^49^ RNA from single cell sorted spike positive memory B cells were reverse transcribed into cDNA for amplification of IgH, Igκ, and Igƛ genes. In brief, cDNA were amplified using two semi-nested PCRs^50^. Sanger sequencing was performed on the PCR products from the second amplification. Sequences were analyzed and aligned using IgBLAST (https://www.ncbi.nlm.nih.gov/igblast/). PCR products were cloned into expression vectors (gifts from Michel C. Nussenzweig, Rockefeller University) by restriction enzyme digestion as previously described^49^. Paired IgH and IgL expression plasmids were transfected using the ExpiFectamine 293 Transfection Kit (ThermoFisher Scientific) according to the manufacturer’s protocol. At seven days post-transfection, cells were centrifuged at 300 g for 5 minutes. Supernatant was collected and purified by protein A (ThermoFisher Scientific). Purified antibody concentrations were measured using NanoDrop (ThermoFisher Scientific) and stored at either 4°C for short term storage or -20°C for long term storage.

### Competition ELISA and antibody epitope grouping

ELISA-based antibody competition assays were performed as previously described. In brief, detection antibodies were biotinylated with EZ-Link Sulfo–NHS-LC-Biotin (ThermoFisher Scientific) following the manufacturer’s protocol. SARS-CoV-2 spike (50 ng/well) were coated onto ELISA plates at 4°C overnight and blocked with 150 μL 4% BSA in PBS for two hours the next day. 30 μL of 2 μg/mL biotinylated antibody was mixed with 30 μL of 200 μg/mL blocking antibody. The mixture was added onto ELISA plates and incubated for 2 hours at 37°C and washed with washing buffer four times. Streptavidin-alkaline phosphatase (50 μL/well; BD Biosciences) was added to the wells following manufacturer’s protocol and incubated at room temperature for one hour. Plates were washed five times with washing buffer and developed at room temperature for two hours. The ELISA plates were read at 405 nm by a microplate reader (Biotek Synergy H1). The detection signal was calculated by (OD value of mixture antibodies-OD value of PBS)/ (OD value of biotinylated antibody alone-OD value of PBS) x100%. Negative values were treated as 100% competition. Competition groups were determined by K-means clustering.

### SARS-CoV-2 pseudovirus neutralization assay

Pseudovirus stocks were produced as described^48^. HEK293T cells were co-transfected with spike encoding (SARS2-spike-delta21-D614G, SARS2-spike-delta21-BA.1, SARS2-spike-delta21-BA.2, SARS2-spike-delta21-BA.5, SARS2-spike-delta21-BQ.1.1, SARS2-spike-delta21-XBB.1.5, SARS2-spike-delta21-EG.5, SARS2-spike-delta21-BA.2.86), package plasmid (psPAX2, Addgene) and backbone plasmid (pLenti CMV Puro LUC, Addgene) with Lipofectamine 3000. Medium was replaced with fresh medium at 24 h, and supernatants were harvested at 48 h post-transfection and clarified by centrifugation at 300 g for 10 min before aliquoting and storing at −80°C. SARS-Cov-2 pseudovirus neutralization assay was performed as described^49^, with target cell line 293FT expressing human ACE2 and serine protease TMPRSS2 (provided by Marc C. Johnson, University of Missouri). Cells at 1.8 × 104 cell/well were seeded in 96-well plates 16 h in advance. Serial diluted mAb was mixed with pseudovirus and incubated for 1 h at 37°C before adding to cells. Cells infected without mAb were scored as 100% infection; cells cultured without pseudovirus or mAb as blank controls. After 48 h incubation at 37°C with 5% CO2, cells were processed with luminescent regent (ONE-Glo™, Promega) according to manufacturer’s instructions, and luminescence (RLU) was measured with a microplate reader (Biotek Synergy H1). Inhibition was calculated by 100-(RLU of mAb-RLU of blank)/ (RLU of pseudovirus -RLU of blank) x100%. Values for half inhibition (IC50) was calculated with GraphPad Prism v10.

### Cryo-EM and negative Staining EM preparation of the G614 Spike–V30V4–SP1-77 Fab Complex

The full-length G614 spike protein containing a C-terminal strep tag was produced following a protocol described previously^51^. To prepare the V30V4-spike complex for cryo-EM analysis, the purified G614 spike was first mixed with SP1-77 Fab at a molar ratio of 1:6, and incubated at room temperature for 30 min. V30V4 Fab was then added to the mixture at a spike-to-Fab molar ratio of 1:6, and incubated for additional 30 min. The antibody-spike complex was further purified by gel-filtration chromatography on a Superose 6 Increase 10/300 column (GE Healthcare) in a buffer containing 25 mM Tris, pH7.5, 150 mM NaCl and 0.02% DDM. To prepare negative staining EM grids, 4 μl of freshly purified peak fraction sample was adsorbed to a glow-discharged carbon-coated copper grid (Electron Microscopy Sciences) and stained with freshly prepared 1.5% uranyl formate. Images were recorded at room temperature on the Tecnai T12 or Phillips CM10 transmission electron microscope with a nominal magnification of 52,000×. Particles were auto-picked and 2D class averages were generated using RELION 4.0.0^52^. The fractions containing the dissociated S1in complex with V30V4 and SP1-77 Fabs were pooled and concentrated.

The concentrated S1-Fab complex was mixed with Bio beads SM2 (Bio-Red) to remove excessive DDM accumulated from the concentrating step. The complex was concentrated again to 7 mg/ml optimal for cryo-grid preparation, and 0.02% DDM was added to the sample to overcome the preferred orientation issue. To prepare cryo-EM grids, 1.2/1.3 Quantifoil Copper grids (Electron Microscopy Sciences), glow-discharged for 60 s at 15 mA using a PELCO easiGlow glow discharge cleaning system (Ted Pella), were coated with the complex and immediately plunge-frozen into liquid ethane using the Vitrobot Mark IV (Thermo Fisher Scientific). Excess protein was blotted away using grade 595 filter paper (Ted Pella) with a blotting time of 5 s, a blotting force of −5 at 4 °C with 100% humidity. The grids were first screened for ice thickness and particle distribution. Selected grids were used to acquire images with a Titan Krios transmission electron microscope (Thermo Fisher Scientific) operated at 300 keV and equipped with a BioQuantum GIF/K3 direct electron detector. Automated data collection was carried out using SerialEM version 3.8.6 with a defocus range of 0.5 to 2.2 μm.

### Cryo-EM data processing

All cryo-EM data processing was carried out using RELION 3.1^53^ and cryoSPARC v4.3.1^54^. The motion correction for the recorded images was performed by patch motion correction, and the contrast transfer function was estimated using CTFFIND4^55^. 577 out of 13,721 images with the CTF fit resolution worse than 5 or severe ice contaminations were excluded manually. Blob Picker was used to select particles for 2D classification and good 2D images were chosen as templates for template-based particle picking. Two different data sets of the G614 Spike S1– V30V4–SP1-77 complex were processed separately with 2D and 3D classification and good particles from both sets were combined for final processing. With Template Picker, total 10,906,583 particles were selected from 13,144 micrographs. After manual inspection of the picked particles and 4-6 rounds of 2D classification, the resulting 1,170,510 particles were subjected to 3-4 rounds of heterogeneous refinement. The initial model was generated by 200,000 particles using ab-initio reconstruction. The best two classes, one from each data set, containing 110,708 and 152,214 particles, respectively, were combined and subjected to non-uniform refinement, giving a map at 2.99 Å. One round Global CTF refinement improved the resolution to 2.95 Å, and to 2.94 Å after additional local refinement with a soft mask covering the rigid G614 Spike–V30V4–SP1-77 interacting domains. To further improve the resolution, Bayesian polishing was performed in RELION with the selected 262,922 particles and raw movies, and the particles were subjected to another round non-uniform refinement in cryoSPARC, leading to a map at 2.94 Å, and at 2.85 Å after further local refinement. A new round of heterogeneous classification removed additional junk particles, producing a map at 2.88 Å with 244,319 particles. Global CTF refinement improved the resolution to 2.85 Å, and further local refinement led to a final map at 2.82 Å for model building.

Reported resolutions were based on the gold-standard Fourier shell correlation (FSC) using the 0.143 criterion. All density maps were corrected for the modulation transfer function (MTF) of the K2 summit direct detector and then sharpened by applying a temperature factor that was estimated by sharping tools in cryoSPARC. Local resolution estimation tool in cryoSPARC was used to determine local resolution of the map.

### Model building

The final map with the correct handedness was used for model building. AlphaFold2^56^ and the tool ModelAngelo building^57^ in RELION were used to predict the initial model of the V30V4 Fab. The SARS-CoV-2 D614G spike-SP1-77 structure (PDB ID: 7UPX) was also used as an initial template for model building. Several rounds of manual building were performed in Coot 0.9.6 ^58^. The model was refined in Phenix 1.20.1-4487 ^59^ against the final map at 2.82 Å. The refinement statistics are summarized in Tabel 1. ChimeraX 1.4 ^60^ was used to visualize cryo-EM maps.

### Protein data bank (PDB) dataset and contact analysis

To investigate structural determinants of antibody recognition of SARS-CoV-2, we analyzed antibody-antigen complexes organized by heavy chain germline genes. 280 PDB files containing antibody-SARS-CoV-2 complexes were processed using a two-stage pipeline to identify relevant chains and organize structures by germline origin for comparative analysis. We identified antibody heavy chain (VH) and light chain (VL) sequences using MUSCLE v5^61^ alignments, with chains showing ≥90% sequence identity to reference sequences classified as antibody chains denoted in CoV-AbDab^39^. Wuhan SARS-CoV-2 RBD chains were identified using three conserved motifs: SNCVADYSVLYNSASFSTF, RGDEVRQIAPGQTGK, and STPCNGVEGFNCYF, with an 80% identity threshold to exclude variant RBDs. For each structure, we extracted only the single complex consisting of VH, VL, and RBD chains, based on minimal Cα-Cα distances. Processed structures were then organized by heavy chain germline genes. Within each germline group, structures were aligned to a reference RBD using PyMOL version 2.1 to standardize orientation for direct comparison of binding modes.

We then analyzed the binding interfaces of 51 class 1 IGHV3-53/66 antibody-SARS-CoV-2 RBD complexes. Antibody-antigen contacts were identified using distance-based criteria, with residues within 4.5Å of partner atoms classified as contact residues using Arpeggio^62^. Then to qualitatively evaluate the epitope sites, RBD residues were classified into three non-exclusive categories: invariant residues (conserved across variants from wild-type to JN.1), variant residues (mutated in at least one major variant), and ACE2-binding footprint residues (involved in ACE2 receptor interaction). We quantified the frequency of antibody contacts for each RBD residue and established multiple frequency thresholds ranging from 90% to 50% of the antibody dataset to define consensus epitopes of varying stringency. This classification was performed separately for heavy and light chain contacts, allowing us to characterize chain-specific contributions to epitope recognition. Then we visualized epitope patterns using ChimeraX to map contact residues onto the RBD structure. To visualize mutation prevalence, we considered the sequences from D614G, BA.1, BA.2, BA.5, and XBB.

### Antigen-specific antibody repertoire datasets and breadth classification

We analyzed antibody gene usage patterns across multiple datasets, including naive B cell repertoires from three studies^63–65^, HIV-1 neutralizing antibodies (CATNAP database^66^), influenza antibodies (Wu Lab^67^), and SARS-CoV-2 antibodies from a centralized database (CoV-AbDab^39^) and a public dataset of neutralization-tested antibodies (Cao^23,29,37,38^). For SARS-CoV-2 antibodies from the Cao dataset, we classified antibodies into the following neutralization categories: WT-only (neutralizes D614G but not Omicron variants), BA.2-broad (neutralizes through BA.2 but not later variants), BA.5-broad (neutralizes through BA.5 but not XBB), and XBB-broad (neutralizes all variants including XBB.1.5). Neutralization was defined as an IC50 ≤ 1 μ g/mL against the respective variant. Heavy chain V gene usage was analyzed across all datasets. Gene names were standardized by removing allele designations, and IGHV3-53 and IGHV3-66 were combined into a single category (IGHV3-53/66) due to their high sequence similarity. We calculated gene frequencies by dividing the count of each gene by the total number of antibodies in each dataset or breadth category. For the naive repertoire analysis, we aggregated gene counts across all donors in the Briney dataset.

### Comparative analysis of heavy chain sequences

We identified all IGHV3-53/66 antibodies in the Cao dataset with available sequencing (179) and filtered for sequences with CDRH3 lengths ≤15 amino acids (156). Sequences were grouped based on two grouping schemes: (1) narrow (only neutralizing Wuhan strain) vs. broad (neutralizing all tested strains through XBB) neutralizers (IC50 ≤ 1 μ g/mL), and (2) Wuhan exposed (from individuals only exposed to Wuhan strain) vs. BTI (from CoronaVac vaccinees with subsequent Omicron infections) antibodies. For antibodies in each group, totals for each amino acid at each position across aligned antibody sequences were calculated. Enrichment significance for amino acid totals at each position between neutralization and exposure groups was assessed by Fisher’s exact test (p < 0.05). The same test was applied to compare sequences in our cohort against BTI sequences from the Cao dataset.

### Generation of average structures of light-chain-defined groups of IGHV3-53/66 antibodies

We filtered all IGHV3-53/66 PDB structures referenced across CoV-AbDab by several criteria: (1) presence of Wuhan RBD, (2) separate chains for VH and VL, (3) CDRH3 length ≤13 amino acids, (4) light chain pairing with either IGKV1-9, IGKV1-33, or IGKV3-20. New structures were created including only chains for VH, VL, and Wuahn RBD. Structures with common light chains were superimposed with respect to RBD in ChimeraX. The average position for each corresponding, common backbone atom (ca, c, n, o) across VHs and VLs within each light-chain-defined group of structures was calculated. Average atomic coordinates were used to generate new “average” structures for IGKV1-9, IGKV1-33, and IGKV3-20 paired IGHV3-53/66 antibodies.

---

**Supplemental Figure 1.**
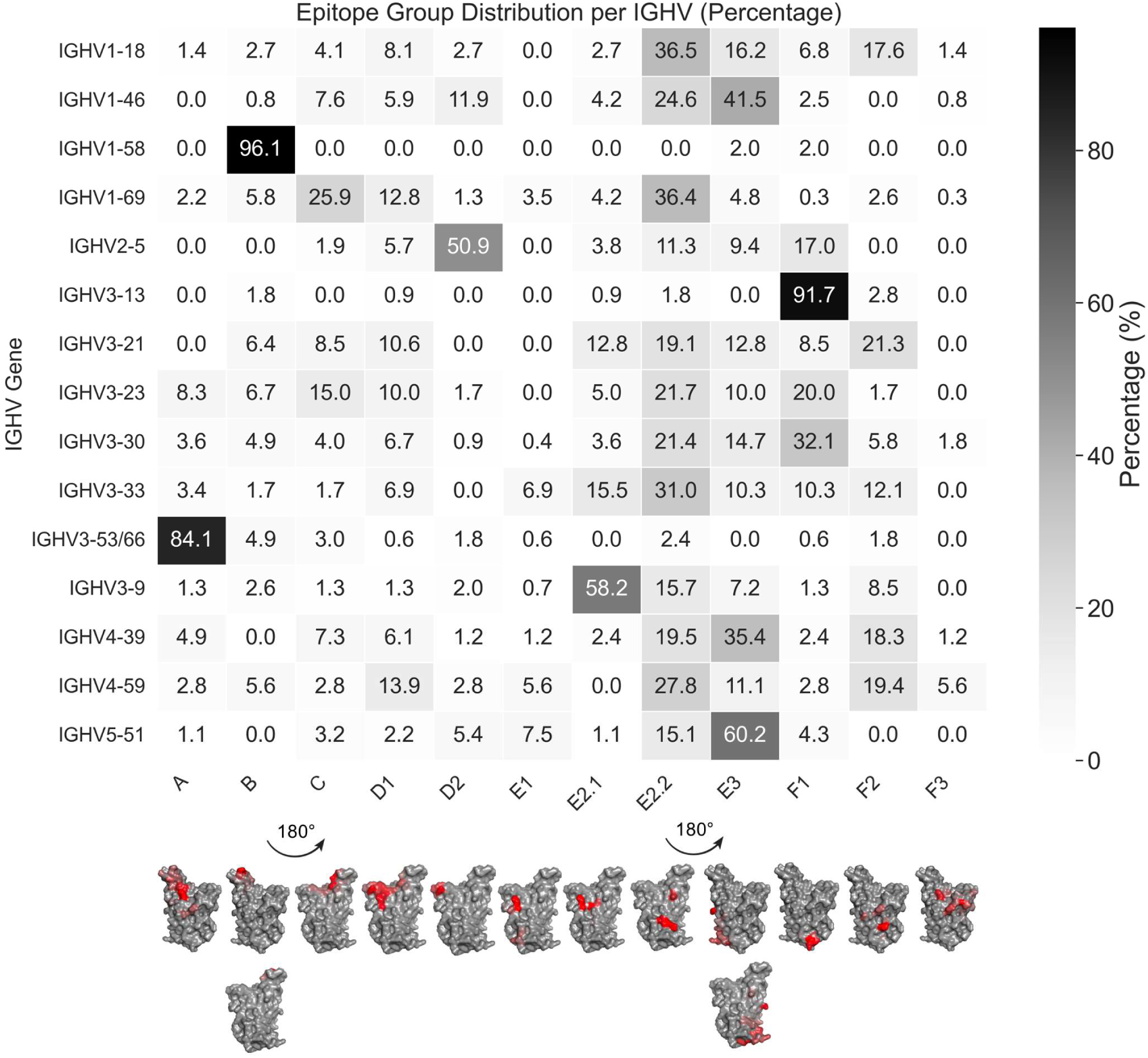
Epitope groups defined by deep mutational scanning profiles (source data: Cao et al., 2023) and their distribution across IGHV genes. Pre-calculated escape scores are mapped onto RBD structures using a red color scale.

**Supplemental Figure 2.**
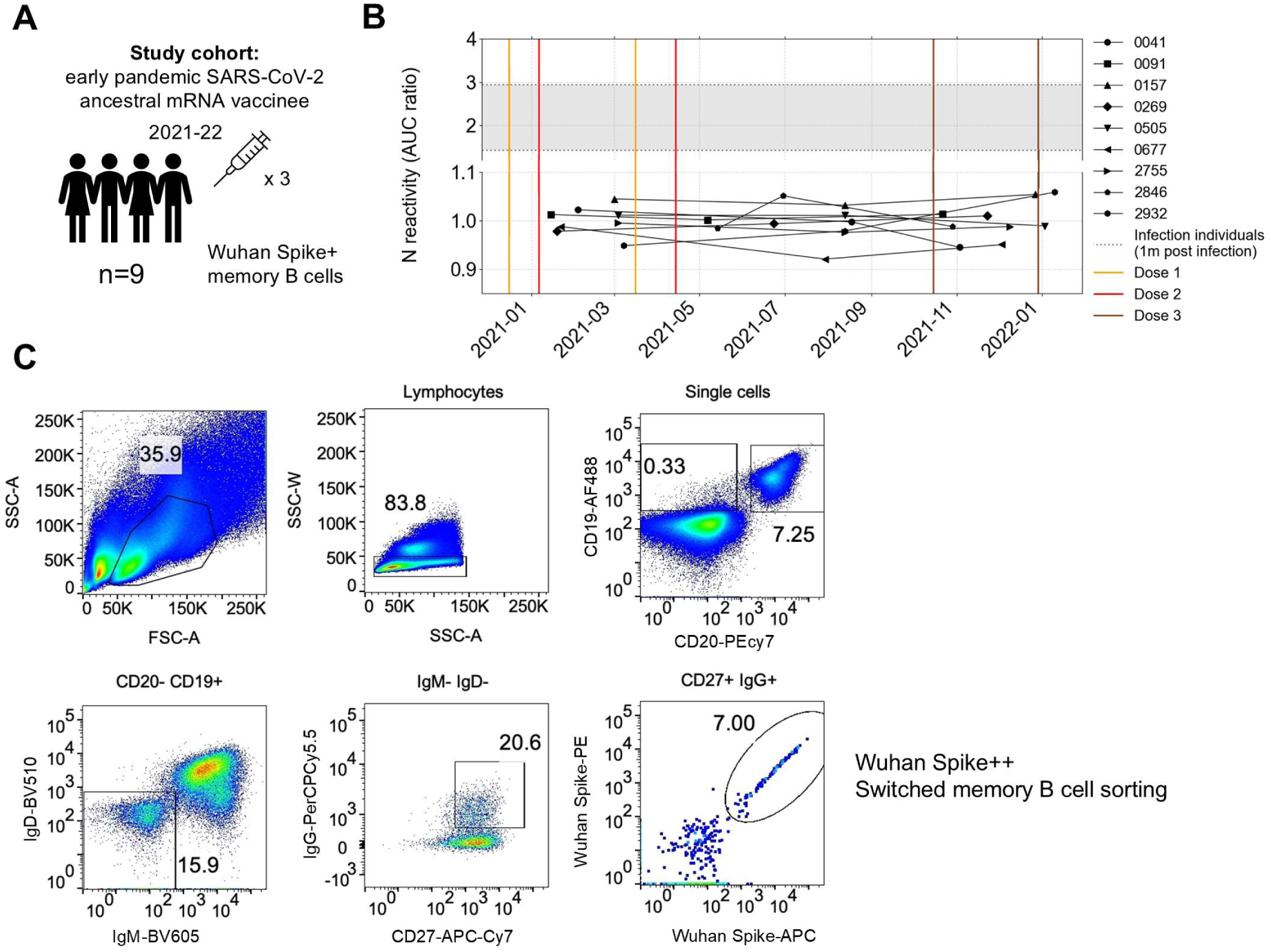
Memory B cell sorting from an infection-naïve, Wuhan-mRNA vaccinee cohort. (A) Cohort summary. (B) Plasma reactivity to nucleocapsid protein from 9 study subjects, and the AUC ratio relative to blank (background control) was plotted. Nine positive control subjects (taken 1 month post-infection) were included. (C) Gating strategy for spike-reactive memory B cell sorting.

**Supplemental Figure 3.**
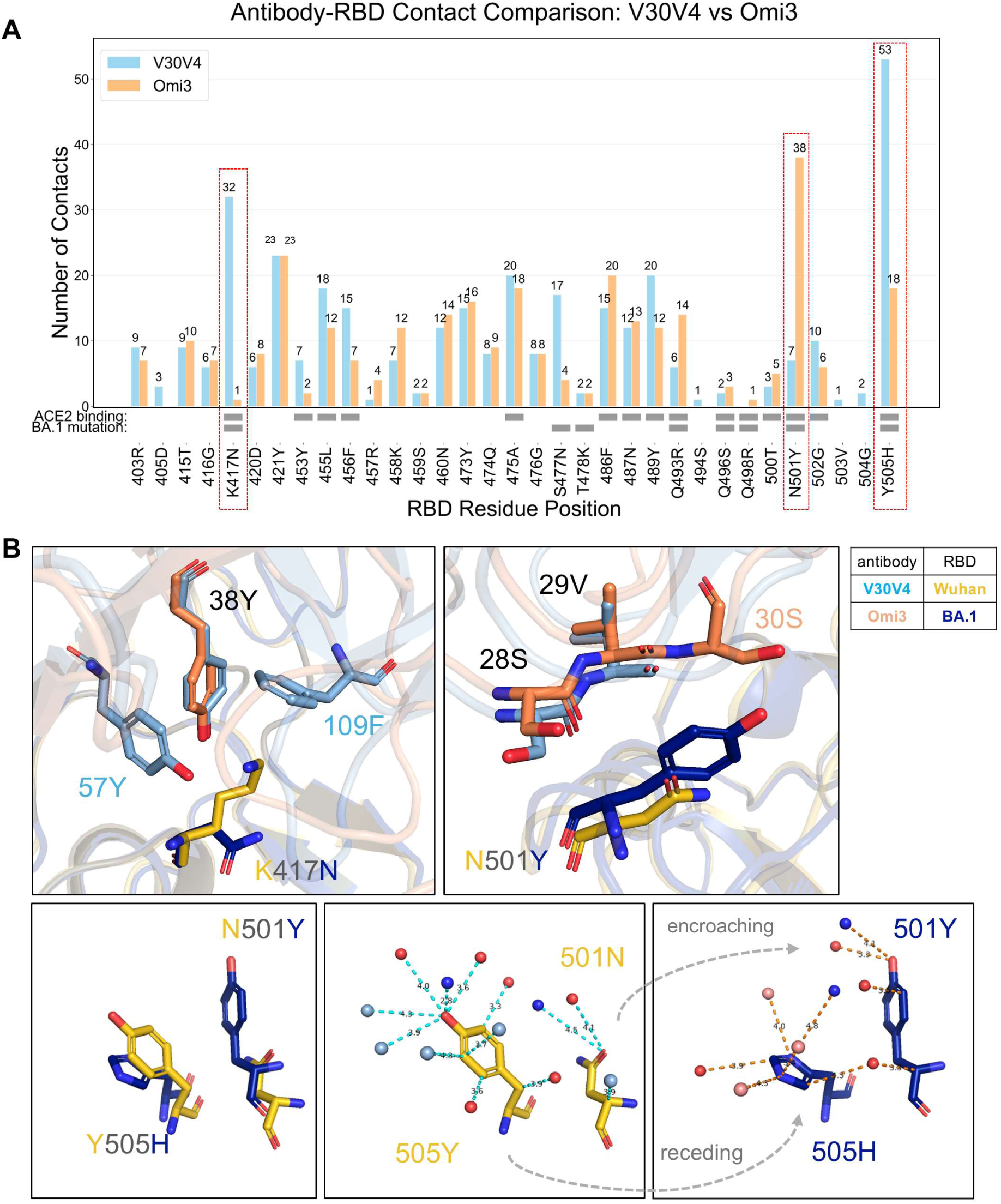
Contact details comparing antibodies V30V4 and Omi3. (A) Atomic contact counts. (B) Representative interaction snapshots showing RBD mutations K417N, N501Y, and Y505H.

**Supplemental Figure 4.**
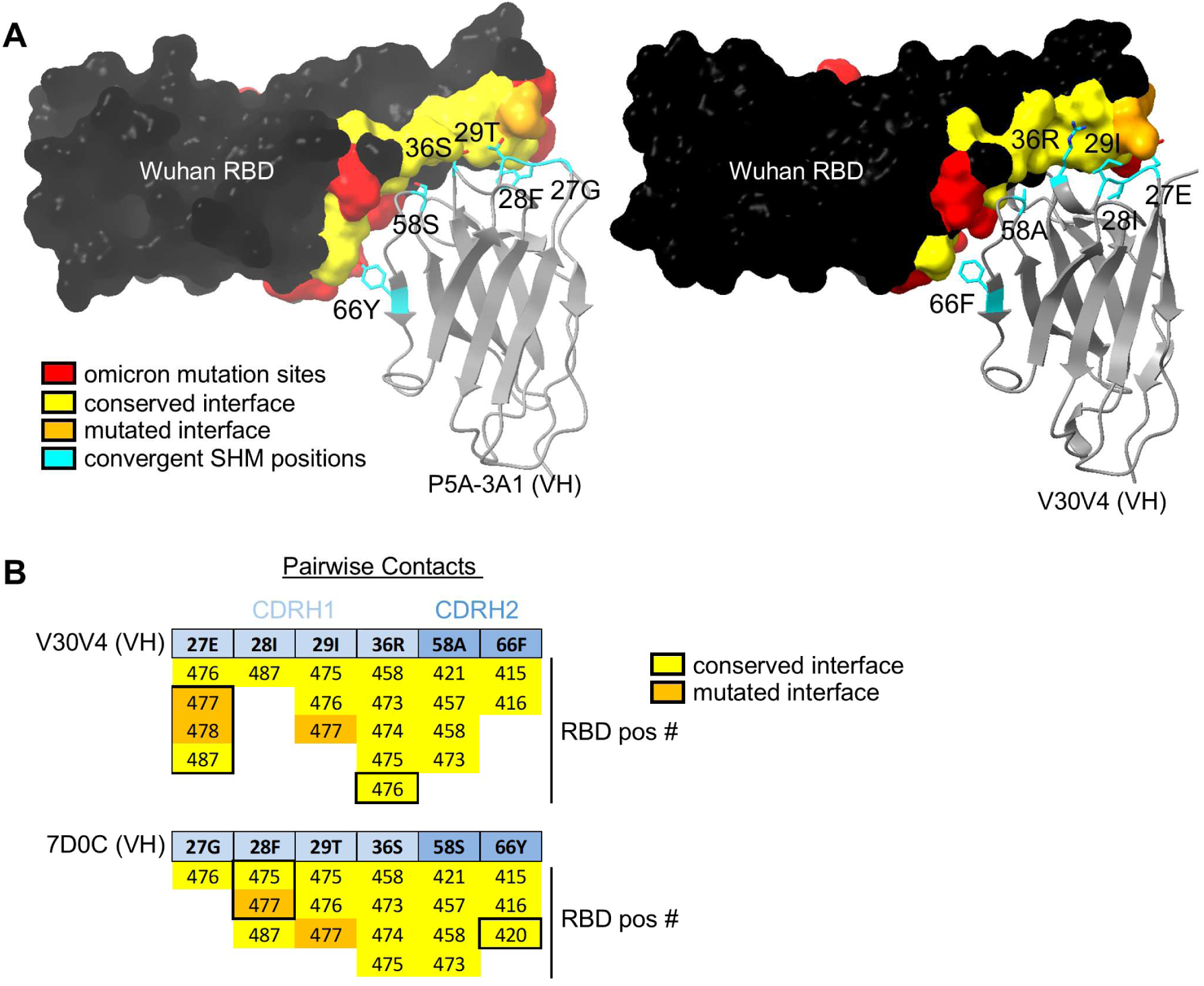
Interface Comparisons between V30V4 and P5A-3A1, a fully-germline IGHV3-53:IGKV3-20 antibody. (A) **I**nteraction surface of Wuhan RBD with both V30V4 and P5A-3A1 (PDB ID: 7D0C) at positions of highly convergent IGHV3-53/66 SHM. (B) Pairwise breakdown of Wuhan RBD interacting positions and hallmark SHM sites on V30V4 and P5A-3A1.

**Supplemental Figure 5.**
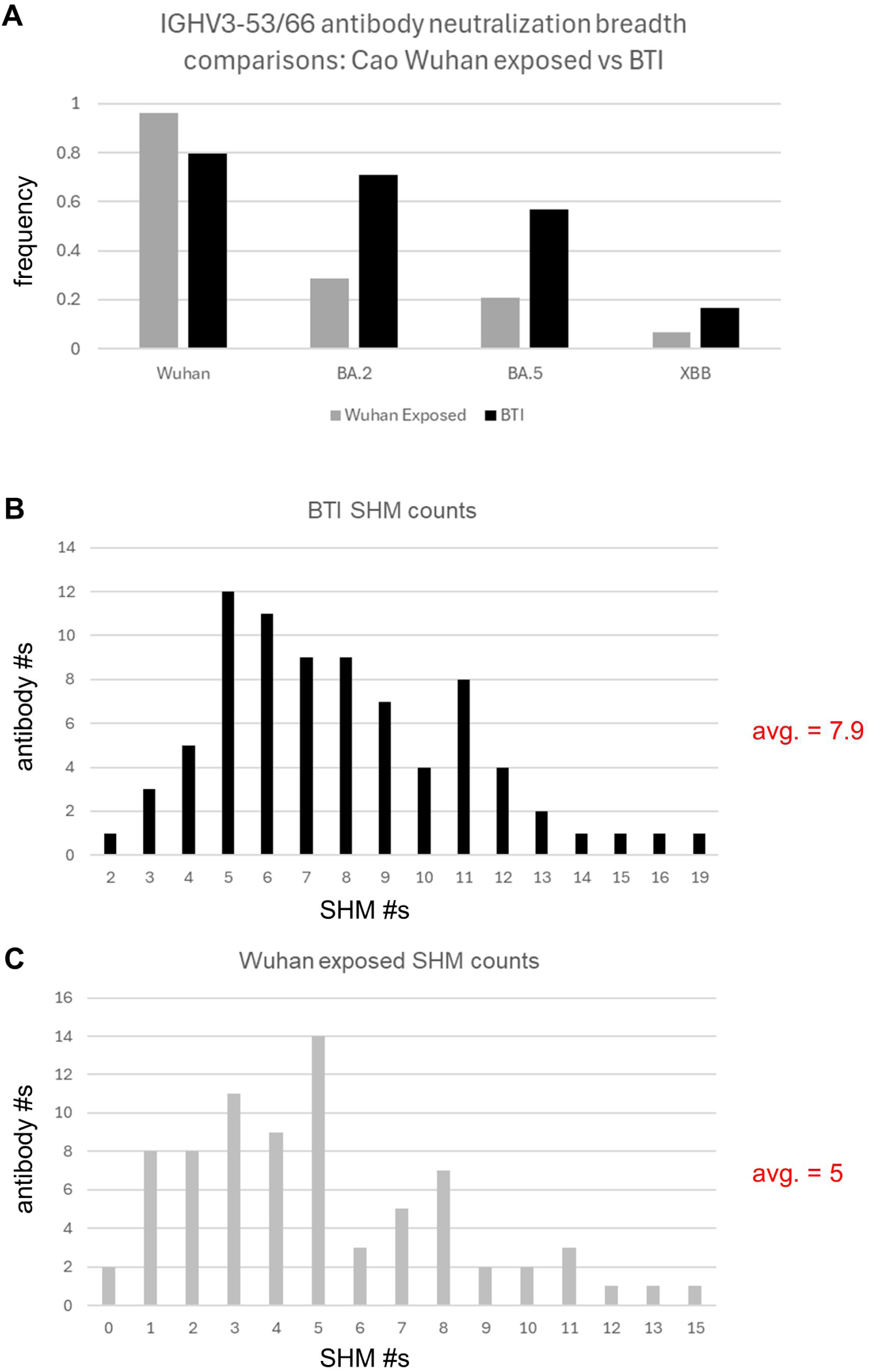
Neutralization Breadth and SHM comparisons between Cao IGHV3-53/66 antibodies by exposure group. (A) Proportion of antibodies in “Wuhan exposed” and “BTI” groups that neutralize all tested strains through the noted strain. (B) SHM counts along VH gene segments for Cao BTI antibodies. (C) SHM counts along VH gene segments for Cao Wuhan exposed antibodies.

**Supplemental Figure 6.**
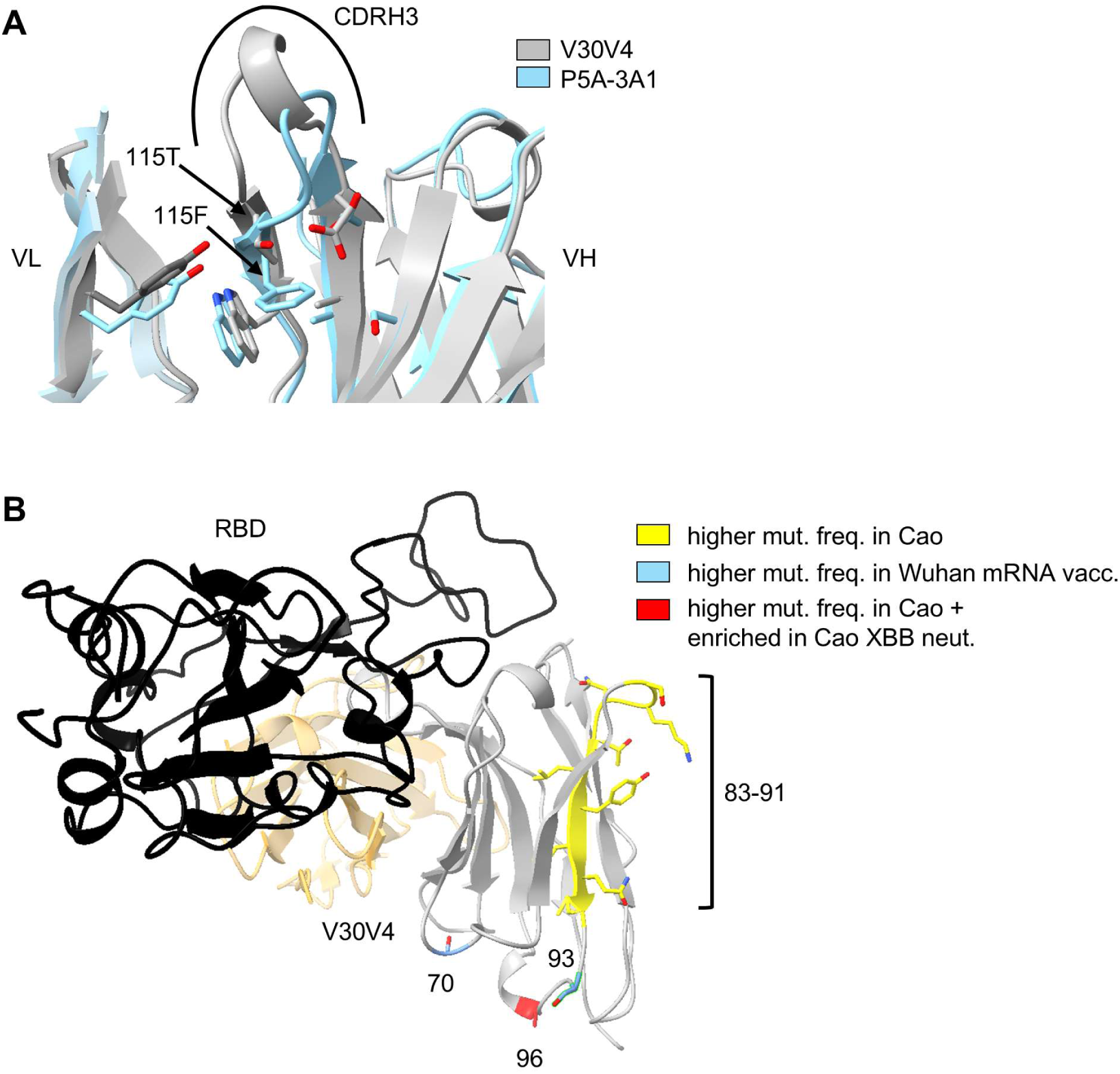
Noteworthy features of IGHV3-53/66 antibodies. (A) Structural comparison between V30V4, containing the highly enriched, neutralization-breadth-associated residue 115T, and P5A-3A1, containing the germline residue 115F. (B) Highlighted positions with differential mutational frequencies between Wuhan-mRNA vaccine derived antibodies from this study, and Cao BTI antibodies.

**Supplemental Figure 7.**
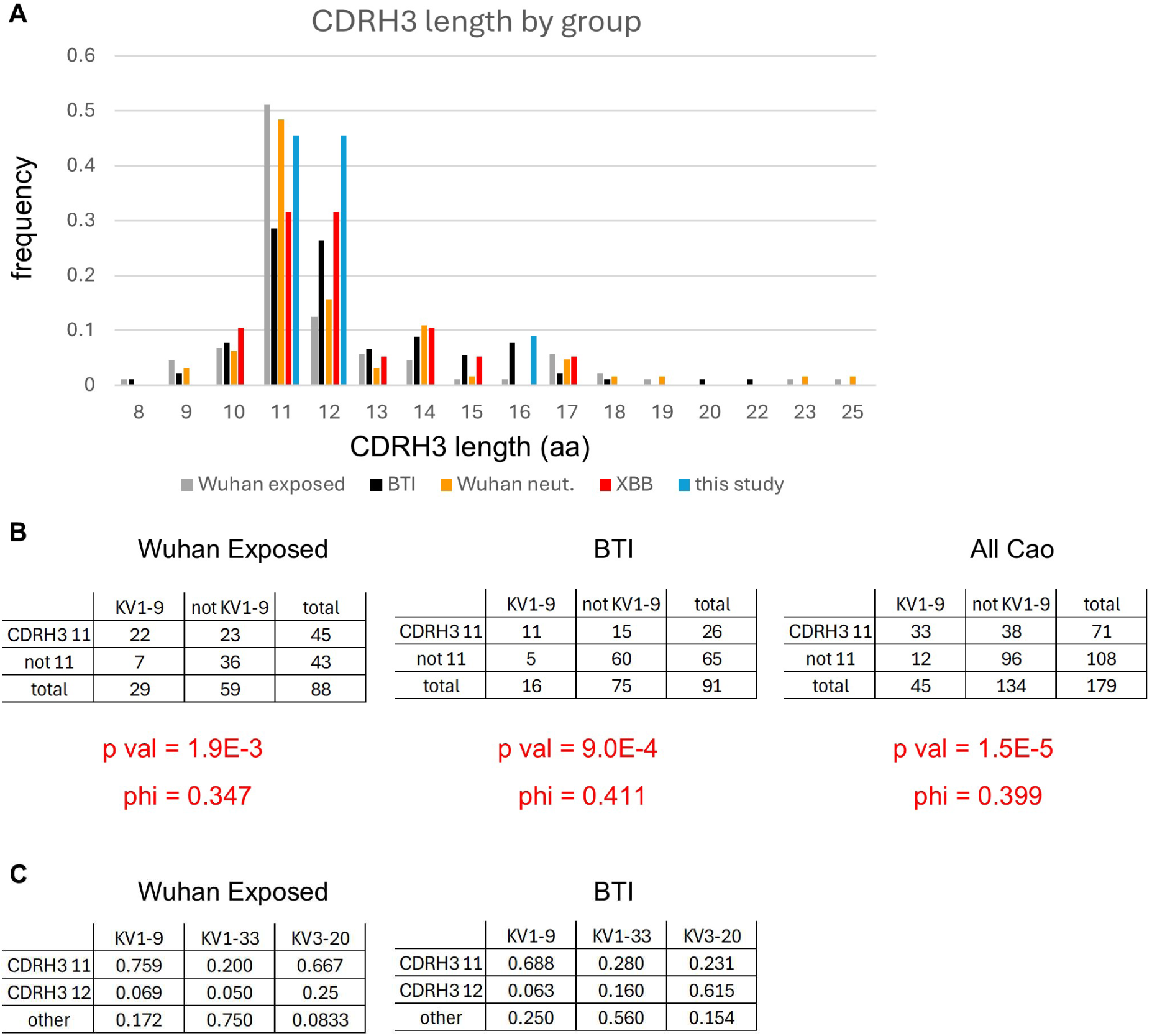
Association between CDRH3 length and light chain pairing. (A) CDRH3 length breakdown for IGHV3-53/66 antibodies by exposure and neutralization breadth category. (B) Counts and statistics for IGHV3-53/66 antibodies with 11aa CDRH3s and IGKV1-9 pairing by exposure group. (C) Overall frequencies of the intersection of CDRH3 lengths and 3 common light chain pairings for IGHV3-53/66 antibodies by exposure group. Data from Cao dataset.

**Supplemental Figure 8.**
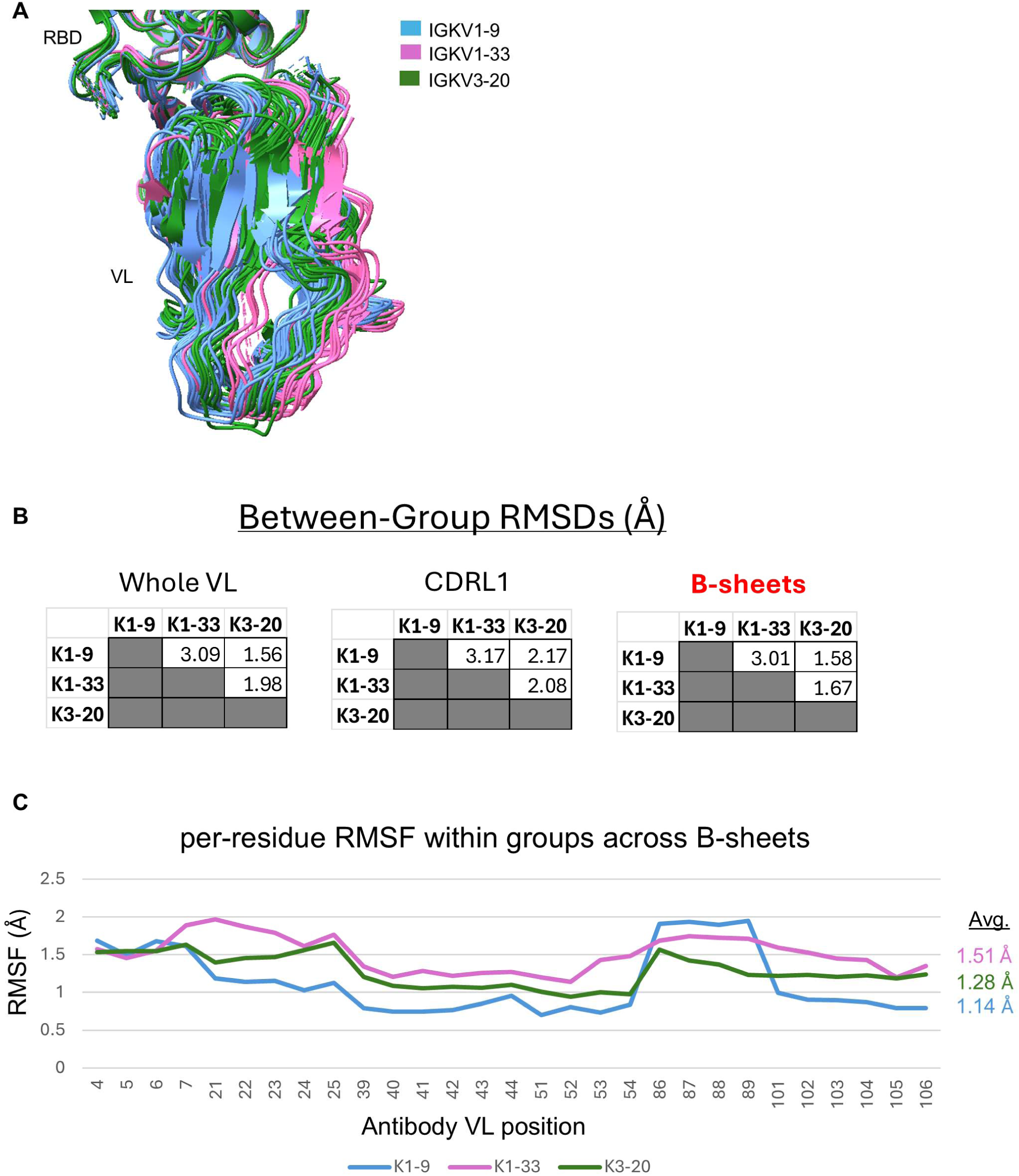
Distance measurements describing positional gradient of IGHV3-53/66 antibodies by paired light chain. (A) Superimposition of all analyzed IGHV3-53/66 antibody structures paired with 3 common light chains. (B) Root mean squared distance (RMSD) calculations between average structures of light-chain defined IGHV3-53/66 antibody groups. Measurements are made across all of VL, across only the CDRL1 region, and across positions constituting conserved B-sheets. (C) Per-residue root mean squared fluctuation (RMSF) measurements across all analyzed IGHV3-53/66 antibody structures paired with 3 common light chains, along positions of conserved B-sheets.

**Supplemental Figure 9.**
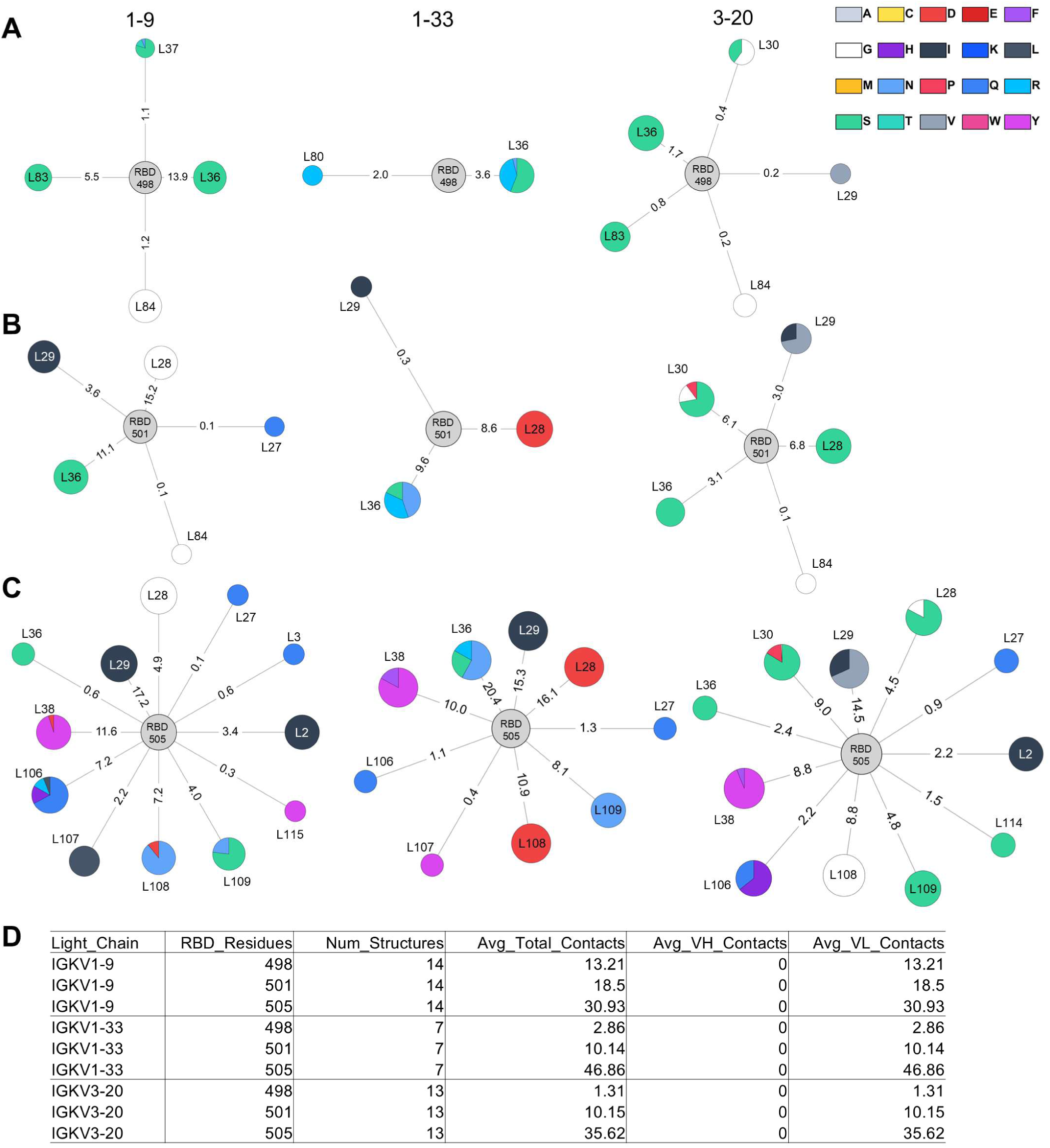
Light chain contact details. Comparing antibody structures paired with IGKV1-9, IGKV1-33, and IGKV3-20 at residues (A) 498, (B) 501, and (C) 505. (D) Average contact frequency per light chain group. In the network diagram, node size reflects the number of contributing PDB structures, while edge length is inversely proportional to the average contact count. Numerical values of average contacts are indicated on the connecting edges.

## References

1. Gaebler, C., Wang, Z., Lorenzi, J.C.C., Muecksch, F., Finkin, S., Tokuyama, M., Cho, A., Jankovic, M., Schaefer-Babajew, D., Oliveira, T.Y., et al. (2021). Evolution of antibody immunity to SARS-CoV-2. Nature 591, 639–644.

2. Gruell, H., Vanshylla, K., Tober-Lau, P., Hillus, D., Schommers, P., Lehmann, C., Kurth, F., Sander, L.E., and Klein, F. (2022). mRNA booster immunization elicits potent neutralizing serum activity against the SARS-CoV-2 Omicron variant. Nat. Med. 28, 477–480.

3. Nachbagauer, R., Wohlbold, T.J., Hirsh, A., Hai, R., Sjursen, H., Palese, P., Cox, R.J., and Krammer, F. (2014). Induction of broadly reactive anti-hemagglutinin stalk antibodies by an H5N1 vaccine in humans. J. Virol. 88, 13260–13268.

4. Wrammert, J., Koutsonanos, D., Li, G.-M., Edupuganti, S., Sui, J., Morrissey, M., McCausland, M., Skountzou, I., Hornig, M., Lipkin, W.I., et al. (2011). Broadly cross-reactive antibodies dominate the human B cell response against 2009 pandemic H1N1 influenza virus infection. J. Exp. Med. 208, 181–193.

5. Ellebedy, A.H., Krammer, F., Li, G.-M., Miller, M.S., Chiu, C., Wrammert, J., Chang, C.Y., Davis, C.W., McCausland, M., Elbein, R., et al. (2014). Induction of broadly cross-reactive antibody responses to the influenza HA stem region following H5N1 vaccination in humans. Proc. Natl. Acad. Sci. U. S. A. 111, 13133–13138.

6. Rusert, P., Kouyos, R.D., Kadelka, C., Ebner, H., Schanz, M., Huber, M., Braun, D.L., Hozé, N., Scherrer, A., Magnus, C., et al. (2016). Determinants of HIV-1 broadly neutralizing antibody induction. Nat. Med. 22, 1260–1267.

7. Guthmiller, J.J., Han, J., Utset, H.A., Li, L., Lan, L.Y.-L., Henry, C., Stamper, C.T., McMahon, M., O’Dell, G., Fernández-Quintero, M.L., et al. (2022). Broadly neutralizing antibodies target a haemagglutinin anchor epitope. Nature 602, 314–320.

8. Whittle, J.R.R., Zhang, R., Khurana, S., King, L.R., Manischewitz, J., Golding, H., Dormitzer, P.R., Haynes, B.F., Walter, E.B., Moody, M.A., et al. (2011). Broadly neutralizing human antibody that recognizes the receptor-binding pocket of influenza virus hemagglutinin. Proc. Natl. Acad. Sci. U. S. A. 108, 14216–14221.

9. Yuan, M., Liu, H., Wu, N.C., Lee, C.-C.D., Zhu, X., Zhao, F., Huang, D., Yu, W., Hua, Y., Tien, H., et al. (2020). Structural basis of a shared antibody response to SARS-CoV-2. Science 369, 1119–1123.

10. Yan, Q., He, P., Huang, X., Luo, K., Zhang, Y., Yi, H., Wang, Q., Li, F., Hou, R., Fan, X., et al. (2021). Germline IGHV3-53-encoded RBD-targeting neutralizing antibodies are commonly present in the antibody repertoires of COVID-19 patients. Emerg. Microbes Infect. 10, 1097–1111.

11. Yan, Q., Zhang, Y., Hou, R., Pan, W., Liang, H., Gao, X., Deng, W., Huang, X., Qu, L., Tang, C., et al. (2024). Deep immunoglobulin repertoire sequencing depicts a comprehensive atlas of spike-specific antibody lineages shared among COVID-19 convalescents. Emerg. Microbes Infect. 13, 2290841.

12. Ge, J., Wang, R., Ju, B., Zhang, Q., Sun, J., Chen, P., Zhang, S., Tian, Y., Shan, S., Cheng, L., et al. (2021). Antibody neutralization of SARS-CoV-2 through ACE2 receptor mimicry. Nat. Commun. 12, 250.

13. Rogers, T.F., Zhao, F., Huang, D., Beutler, N., Burns, A., He, W.-T., Limbo, O., Smith, C., Song, G., Woehl, J., et al. (2020). Isolation of potent SARS-CoV-2 neutralizing antibodies and protection from disease in a small animal model. Science 369, 956–963.

14. Kim, S.I., Noh, J., Kim, S., Choi, Y., Yoo, D.K., Lee, Y., Lee, H., Jung, J., Kang, C.K., Song, K.-H., et al. (2021). Stereotypic neutralizing VH antibodies against SARS-CoV-2 spike protein receptor binding domain in patients with COVID-19 and healthy individuals. Sci. Transl. Med. 13, eabd6990.

15. Gruell, H., Vanshylla, K., Weber, T., Barnes, C.O., Kreer, C., and Klein, F. (2022). Antibody-mediated neutralization of SARS-CoV-2. Immunity 55, 925–944.

16. Shu, Y., and McCauley, J. (2017). GISAID: Global initiative on sharing all influenza data - from vision to reality. Euro Surveill. 22. 10.2807/1560-7917.ES.2017.22.13.30494.

17. Nutalai, R., Zhou, D., Tuekprakhon, A., Ginn, H.M., Supasa, P., Liu, C., Huo, J., Mentzer, A.J., Duyvesteyn, H.M.E., Dijokaite-Guraliuc, A., et al. (2022). Potent cross-reactive antibodies following Omicron breakthrough in vaccinees. Cell 185, 2116–2131.e18.

18. Bruhn, M., Obara, M., Salam, A., Costa, B., Ziegler, A., Waltl, I., Pavlou, A., Hoffmann, M., Graalmann, T., Pöhlmann, S., et al. (2024). Diversification of the VH3-53 immunoglobulin gene segment by somatic hypermutation results in neutralization of SARS-CoV-2 virus variants. Eur. J. Immunol. 54, e2451056.

19. Windsor, I.W., Tong, P., Lavidor, O., Moghaddam, A.S., McKay, L.G.A., Gautam, A., Chen, Y., MacDonald, E.A., Yoo, D.K., Griffths, A., et al. (2022). Antibodies induced by an ancestral SARS-CoV-2 strain that cross-neutralize variants from Alpha to Omicron BA.1. Sci. Immunol. 7, eabo3425.

20. Paciello, I., Maccari, G., Pierleoni, G., Perrone, F., Realini, G., Troisi, M., Anichini, G., Cusi, M.G., Rappuoli, R., and Andreano, E. (2024). SARS-CoV-2 JN.1 variant evasion of IGHV3-53/3-66 B cell germlines. Sci. Immunol. 9, eadp9279.

21. Luo, M., Zhou, R., Tang, B., Liu, H., Chen, B., Liu, N., Mo, Y., Zhang, P., Lee, Y.L., Ip, J.D., et al. (2024). Ultrapotent class I neutralizing antibodies post Omicron breakthrough infection overcome broad SARS-CoV-2 escape variants. EBioMedicine 108, 105354.

22. Kuwata, T., Kaku, Y., Biswas, S., Matsumoto, K., Shimizu, M., Kawanami, Y., Uraki, R., Okazaki, K., Minami, R., Nagasaki, Y., et al. (2024). Induction of IGHV3-53 public antibodies with broadly neutralising activity against SARS-CoV-2 including Omicron subvariants in a Delta breakthrough infection case. EBioMedicine 110, 105439.

23. Jian, F., Wang, J., Yisimayi, A., Song, W., Xu, Y., Chen, X., Niu, X., Yang, S., Yu, Y., Wang, P., et al. (2025). Evolving antibody response to SARS-CoV-2 antigenic shift from XBB to JN.1. Nature 637, 921–929.

24. Muecksch, F., Wang, Z., Cho, A., Gaebler, C., Ben Tanfous, T., DaSilva, J., Bednarski, E., Ramos, V., Zong, S., Johnson, B., et al. (2022). Increased memory B cell potency and breadth after a SARS-CoV-2 mRNA boost. Nature 607, 128–134.

25. Li, L., Chen, X., Wang, Z., Li, Y., Wang, C., Jiang, L., and Zuo, T. (2023). Breakthrough infection elicits hypermutated IGHV3-53/3-66 public antibodies with broad and potent neutralizing activity against SARS-CoV-2 variants including the emerging EG.5 lineages. PLoS Pathog. 19, e1011856.

26. Malladi, S.K., Jaiswal, D., Ying, B., Alsoussi, W.B., Darling, T.L., Dadonaite, B., Civljak, A., Horvath, S.C., Zhou, J.Q., Kim, W., et al. (2024). Defining a highly conserved B cell epitope in the receptor binding motif of SARS-CoV-2 spike glycoprotein. bioRxivorg. 10.1101/2024.12.06.625234.

27. Hastie, K.M., Yu, X., Ana-Sosa-Batiz, F., Zyla, D.S., Harkins, S.S., Hariharan, C., Wasserman, H., Zandonatti, M.A., Miller, R., Maule, E., et al. (2023). Potent Omicron-neutralizing antibodies isolated from a patient vaccinated 6 months before Omicron emergence. Cell Rep. 42, 112421.

28. Muecksch, F., Weisblum, Y., Barnes, C.O., Schmidt, F., Schaefer-Babajew, D., Wang, Z., C Lorenzi, J.C., Flyak, A.I., DeLaitsch, A.T., Huey-Tubman, K.E., et al. (2021). Affinity maturation of SARS-CoV-2 neutralizing antibodies confers potency, breadth, and resilience to viral escape mutations. Immunity 54, 1853–1868.e7.

29. Cao, Y., Jian, F., Wang, J., Yu, Y., Song, W., Yisimayi, A., Wang, J., An, R., Chen, X., Zhang, N., et al. (2023). Imprinted SARS-CoV-2 humoral immunity induces convergent Omicron RBD evolution. Nature 614, 521–529.

30. Sheward, D.J., Pushparaj, P., Das, H., Greaney, A.J., Kim, C., Kim, S., Hanke, L., Hyllner, E., Dyrdak, R., Lee, J., et al. (2024). Structural basis of broad SARS-CoV-2 cross-neutralization by affinity-matured public antibodies. Cell Rep. Med. 5, 101577.

31. Park, S., Choi, J., Lee, Y., Noh, J., Kim, N., Lee, J., Cho, G., Kim, S., Yoo, D.K., Kang, C.K., et al. (2024). An ancestral SARS-CoV-2 vaccine induces anti-Omicron variants antibodies by hypermutation. Nat. Commun. 15, 3368.

32. Weber, T., Dähling, S., Rose, S., Affeldt, P., Vanshylla, K., Ullrich, L., Gieselmann, L., Teipel, F., Gruell, H., Di Cristanziano, V., et al. (2023). Enhanced SARS-CoV-2 humoral immunity following breakthrough infection builds upon the preexisting memory B cell pool. Sci. Immunol. 8, eadk5845.

33. Garcia-Beltran, W.F., St Denis, K.J., Hoelzemer, A., Lam, E.C., Nitido, A.D., Sheehan, M.L., Berrios, C., Ofoman, O., Chang, C.C., Hauser, B.M., et al. (2022). mRNA-based COVID-19 vaccine boosters induce neutralizing immunity against SARS-CoV-2 Omicron variant. Cell 185, 457–466.e4.

34. Nemet, I., Kliker, L., Lustig, Y., Zuckerman, N., Erster, O., Cohen, C., Kreiss, Y., Alroy-Preis, S., Regev-Yochay, G., Mendelson, E., et al. (2022). Third BNT162b2 vaccination neutralization of SARS-CoV-2 omicron infection. N. Engl. J. Med. 386, 492–494.

35. Muik, A., Lui, B.G., Wallisch, A.-K., Bacher, M., Mühl, J., Reinholz, J., Ozhelvaci, O., Beckmann, N., Güimil Garcia, R. de la C., Poran, A., et al. (2022). Neutralization of SARS-CoV-2 Omicron by BNT162b2 mRNA vaccine-elicited human sera. Science 375, 678–680.

36. Liang, C.-Y., Raju, S., Liu, Z., Li, Y., Asthagiri Arunkumar, G., Case, J.B., Scheaffer, S.M., Zost, S.J., Acreman, C.M., Gagne, M., et al. (2024). Imprinting of serum neutralizing antibodies by Wuhan-1 mRNA vaccines. Nature 630, 950–960.

37. Cao, Y., Yisimayi, A., Jian, F., Song, W., Xiao, T., Wang, L., Du, S., Wang, J., Li, Q., Chen, X., et al. (2022). BA.2.12.1, BA.4 and BA.5 escape antibodies elicited by Omicron infection. Nature 608, 593–602.

38. Yisimayi, A., Song, W., Wang, J., Jian, F., Yu, Y., Chen, X., Xu, Y., Yang, S., Niu, X., Xiao, T., et al. (2024). Repeated Omicron exposures override ancestral SARS-CoV-2 immune imprinting. Nature 625, 148–156.

39. Raybould, M.I.J., Kovaltsuk, A., Marks, C., and Deane, C.M. (2021). CoV-AbDab: the coronavirus antibody database. Bioinformatics 37, 734–735.

40. Tan, T.J.C., Yuan, M., Kuzelka, K., Padron, G.C., Beal, J.R., Chen, X., Wang, Y., Rivera-Cardona, J., Zhu, X., Stadtmueller, B.M., et al. (2021). Sequence signatures of two public antibody clonotypes that bind SARS-CoV-2 receptor binding domain. Nat. Commun. 12, 3815.

41. Tian, X., Zhu, X., Song, W., Yang, Z., Wu, Y., and Ying, T. (2022). The prominent role of a CDR1 somatic hypermutation for convergent IGHV3-53/3-66 antibodies in binding to SARS-CoV-2. Emerg. Microbes Infect. 11, 1186–1190.

42. Wang, M., Fan, Q., Zhou, B., Ye, H., Shen, S., Yu, J., Cheng, L., Ge, X., Ju, B., and Zhang, Z. (2022). A key F27I substitution within HCDR1 facilitates the rapid maturation of P2C-1F11-like neutralizing antibodies in a SARS-CoV-2-infected donor. Cell Rep. 40, 111335.

43. Zhang, Q., Ju, B., Ge, J., Chan, J.F.-W., Cheng, L., Wang, R., Huang, W., Fang, M., Chen, P., Zhou, B., et al. (2021). Potent and protective IGHV3-53/3-66 public antibodies and their shared escape mutant on the spike of SARS-CoV-2. Nat. Commun. 12, 4210.

44. Lefranc, M.-P., Pommié, C., Ruiz, M., Giudicelli, V., Foulquier, E., Truong, L., Thouvenin-Contet, V., and Lefranc, G. (2003). IMGT unique numbering for immunoglobulin and T cell receptor variable domains and Ig superfamily V-like domains. Dev. Comp. Immunol. 27, 55– 77.

45. Wu, N.C., Yuan, M., Liu, H., Lee, C.-C.D., Zhu, X., Bangaru, S., Torres, J.L., Caniels, T.G., Brouwer, P.J.M., van Gils, M.J., et al. (2020). An alternative binding mode of IGHV3-53 antibodies to the SARS-CoV-2 receptor binding domain. Cell Rep. 33, 108274.

46. Korenkov, M., Zehner, M., Cohen-Dvashi, H., Borenstein-Katz, A., Kottege, L., Janicki, H., Vanshylla, K., Weber, T., Gruell, H., Koch, M., et al. (2023). Somatic hypermutation introduces bystander mutations that prepare SARS-CoV-2 antibodies for emerging variants. Immunity 56, 2803–2815.e6.

47. Francis, T. (1960). On the Doctrine of Orignal Antigenic Sin. Proc. Am. Philos. Soc. 104, 572–578

48. Chen, Y., Zuiani, A., Fischinger, S., Mullur, J., Atyeo, C., Travers, M., Lelis, F.J.N., Pullen, K.M., Martin, H., Tong, P., et al. (2020). Quick COVID-19 healers sustain anti-SARS-CoV-2 antibody production. Cell 183, 1496–1507.e16.

49. Tong, P., Gautam, A., Windsor, I.W., Travers, M., Chen, Y., Garcia, N., Whiteman, N.B., McKay, L.G.A., Storm, N., Malsick, L.E., et al. (2021). Memory B cell repertoire for recognition of evolving SARS-CoV-2 spike. Cell 184, 4969–4980.e15.

50. Tiller, T., Meffre, E., Yurasov, S., Tsuiji, M., Nussenzweig, M.C., and Wardemann, H. (2008). Efficient generation of monoclonal antibodies from single human B cells by single cell RT-PCR and expression vector cloning. J. Immunol. Methods 329, 112–124.

51. Kibria, M.G., Lavine, C.L., Tang, W., Wang, S., Gao, H., Shi, W., Zhu, H., Voyer, J., Rits-Volloch, S., Keerti, et al. (2023). Antibody-mediated SARS-CoV-2 entry in cultured cells. EMBO Rep. 24, e57724.

52. Zivanov, J., Nakane, T., Forsberg, B.O., Kimanius, D., Hagen, W.J.H., Lindahl, E., and Scheres, S.H.W. (2022). New tools for automated cryo-EM single-particle analysis in RELION-4.0. Biophys. J. 121, 2572–2585.

53. Zivanov, J., Nakane, T., and Scheres, S.H.W. (2018). RELION-3: new tools for automated high-resolution cryo-EM structure determination. Elife 7, e42166.

54. Punjani, A., Rubinstein, J. L., Fleet, D. J. & Brubaker, M. A. (2017). cryoSPARC: algorithms for rapid unsupervised cryo-EM structure determination. Nat. Methods 14, 290–296.

55. Rohou, A. and Grigorieff, N. (2015). CTFFIND4: Fast and accurate defocus estimation from electron micrographs. Journal of structural biology, 192(2), pp.216–221.

56. Jumper, J., Evans, R., Pritzel, A. et al.(2021). Highly accurate protein structure prediction with AlphaFold. Nature 596, 583–589.

57. Jamali, K., Käll, L., Zhang, R. et al. (2024). Automated model building and protein identification in cryo-EM maps. Nature 628, 450–457.

58. Emsley, P., Lohkamp, B., Scott, W.G. and Cowtan, K. (2010). Features and development of Coot. Biological crystallography, 66(4), pp.486–501.

59. Adams, P.D., Afonine, P.V., Bunkóczi, G., Chen, V.B., Davis, I.W., Echols, N., Headd, J.J., Hung, L.W., Kapral, G.J., Grosse-Kunstleve, R.W. and McCoy, A.J. (2010). PHENIX: a comprehensive Python-based system for macromolecular structure solution. Biological crystallography, 66(2), pp.213–221.

60. Pettersen, E.F., Goddard, T.D., Huang, C.C., Meng, E.C., Couch, G.S., Croll, T.I., Morris, J.H., and Ferrin, T.E. (2021). UCSF ChimeraX: Structure visualization for researchers, educators, and developers. Protein Sci. 30, 70–82.

61. Edgar, R.C. (2004). MUSCLE: multiple sequence alignment with high accuracy and high throughput. Nucleic Acids Res. 32, 1792–1797.

62. Jubb, H.C., Higueruelo, A.P., Ochoa-Montaño, B., Pitt, W.R., Ascher, D.B., and Blundell, T.L. (2017). Arpeggio: A web server for calculating and visualising interatomic interactions in protein structures. J. Mol. Biol. 429, 365–371.

63. Briney, B., Inderbitzin, A., Joyce, C., and Burton, D.R. (2019). Commonality despite exceptional diversity in the baseline human antibody repertoire. Nature 566, 393–397.

64. Soto, C., Bombardi, R.G., Branchizio, A., Kose, N., Matta, P., Sevy, A.M., Sinkovits, R.S., Gilchuk, P., Finn, J.A., and Crowe, J.E., Jr (2019). High frequency of shared clonotypes in human B cell receptor repertoires. Nature 566, 398–402.

65. Ghraichy, M., Galson, J.D., Kovaltsuk, A., von Niederhäusern, V., Pachlopnik Schmid, J., Recher, M., Jauch, A.J., Miho, E., Kelly, D.F., Deane, C.M., et al. (2020). Maturation of the human immunoglobulin heavy chain repertoire with age. Front. Immunol. 11, 1734.

66. Yoon, H., Macke, J., West, A.P., Jr, Foley, B., Bjorkman, P.J., Korber, B., and Yusim, K. (2015). CATNAP: a tool to compile, analyze and tally neutralizing antibody panels. Nucleic Acids Res. 43, W213–9.

67. Wang, Y., Lv, H., Teo, Q.W., Lei, R., Gopal, A.B., Ouyang, W.O., Yeung, Y.-H., Tan, T.J.C., Choi, D., Shen, I.R., et al. (2024). An explainable language model for antibody specificity prediction using curated influenza hemagglutinin antibodies. Immunity 57, 2453–2465.e7.

